# Regulation of the poly(A) Polymerase Star-PAP by a Nuclear Phosphoinositide Signalosome

**DOI:** 10.1101/2024.07.01.601467

**Authors:** Tianmu Wen, Mo Chen, Vincent L. Cryns, Richard A. Anderson

**Affiliations:** University of Wisconsin Carbone Cancer Center, University of Wisconsin-Madison, School of Medicine and Public Health; 1111 Highland Avenue, Madison, WI 53705, USA; Department of Medicine, University of Wisconsin-Madison, School of Medicine and Public Health; 1111 Highland Avenue, Madison, WI 53705, USA; Current Address: Department of Pharmacology, Joint Laboratory of Guangdong-Hong Kong Universities for Vascular Homeostasis and Diseases, School of Medicine, Southern University of Science and Technology, Shenzhen 518055, China; These authors contributed equally to this work

**Keywords:** Phosphatidylinositol transfer proteins, Phosphoinositide, Phosphoinositide kinases, PIP_n_-linked proteins, Small heat shock proteins, Star-PAP

## Abstract

Star-PAP is a noncanonical poly(A) polymerase that controls gene expression. Star-PAP was previously reported to bind the phosphatidylinositol 4-phosphate 5-kinase PIPKI⍺ and its product phosphatidylinositol 4,5-bisphosphate, which regulate Star-PAP poly(A) polymerase activity and expression of specific genes. Recent studies have revealed a nuclear PI signaling pathway in which the PI transfer proteins PITP⍺/β, PI kinases and phosphatases bind p53 to sequentially modify protein-linked phosphatidylinositol phosphates and regulate its function. Here we demonstrate that multiple phosphoinositides, including phosphatidylinositol 4-monophosphate and phosphatidylinositol 3,4,5-trisphosphate are also coupled to Star-PAP in response to stress. This is initiated by PITP⍺/β binding to Star-PAP, while the Star-PAP-linked phosphoinositides are modified by PI4KII⍺, PIPKI⍺, IPMK, and PTEN recruited to Star-PAP. The phosphoinositide coupling enhances the association of the small heat shock proteins HSP27/⍺B-crystallin with Star-PAP. Knockdown of the PITPs, kinases, or HSP27 reduce the expression of Star-PAP targets. Our results demonstrate that the PITPs generate Star-PAP-PIPn complexes that are then modified by PI kinases/phosphatases and small heat shock proteins that regulate the linked phosphoinositide phosphorylation and Star-PAP activity in response to stress.

## Introduction

Phosphoinositides (PIPs) are a family of minor phospholipids in membranes originating from phosphatidylinositol that have well established roles as second messengers in receptor signaling and membrane trafficking(Balla, 2006; Falkenburger *et al*, 2010; Hammond & Burke, 2020; Sasaki *et al*, 2009). PIs and their associated PI kinases and phosphatases are also present in membrane-free regions of the nucleus where their functional roles are emerging(Boronenkov *et al*, 1998; Chen *et al*, 2020; Cocco *et al*, 1987; Jacobsen *et al*, 2019; Smith & Wells, 1983).

The speckle-targeted PIPKIα regulated-poly(A) polymerase (Star-PAP/TUT1) was the first nuclear enzyme reported to bind PIPs (Mellman *et al*, 2008). Star-PAP is a terminal nucleotidyltransferase that controls the expression of approximately 40% of human genes through alternative polyadenylation (Li *et al*, 2017; Sudheesh *et al*, 2019) and U6 snRNA oligouridylation (Trippe *et al*, 2006; Trippe *et al*, 2003; Trippe *et al*, 1998; Yamashita & Tomita, 2023). These genes have been implicated in diverse cellular processes including oxidative, genotoxic stress and many others (Gonzales *et al*, 2008; Laishram & Anderson, 2010; Laishram *et al*, 2011; Li *et al*, 2012; Li *et al*., 2017; Mellman *et al*., 2008). Star-PAP processes mRNAs by directly binding to the 3’ UTR upstream of the poly(A) signal and recruits cleavage and polyadenylation specificity factor (CPSF) subunits (Laishram & Anderson, 2010). Under some conditions Star-PAP also exhibits terminal uridylyl transferase (TUT1) activity (Trippe *et al*., 2006); however, its preference for ATP over UTP indicates that Star-PAP is predominantly a poly (A) polymerase (PAP)(Mellman *et al*., 2008). Notably, Star-PAP binds the PIP 5-kinase PIPKI⍺ (PIP5K1A) and its product phosphatidylinositol 4,5-bisphosphate (PI(4,5)P_2_), which regulates its PAP activity and controls expression of Star-PAP target mRNAs (Barlow *et al*, 2010; Mellman *et al*., 2008)., including HO-1 (Heme oxygenase-1), BIK (Bcl-2-Interacting Killer), cadherin 1, and cadherin 13 (A P & Laishram, 2018; Koshre *et al*, 2021; Laishram & Anderson, 2010; Laishram *et al*., 2011; Li & Anderson, 2014; Li *et al*, 2013; Li *et al*., 2012; Li *et al*., 2017; Mellman & Anderson, 2009; Mellman *et al*., 2008). Oxidative stress induces Star-PAP phosphorylation by casein kinase I (CKI), which is critical for its PAP activity and priming required for PI(4,5)P_2_ stimulation, leading to HO-1 mRNA processing (Laishram *et al*., 2011). In response to genotoxic stress, nuclear PKCδ is recruited to the Star-PAP complex and the bound PI(4,5)P_2_ on Star-PAP activates PKCδ, which phosphorylates Star-PAP, regulating the processing of BIK mRNA (Li *et al*., 2012). Hence, PI(4,5)P_2_ binding to Star-PAP plays a critical role in controlling its PAP activity and mRNA regulation.

Recent studies have revealed a nuclear PI signaling pathway in which the PI transfer proteins PITP⍺/β, PI kinases and phosphatases bind p53 to sequentially modify phosphatidylinositol phosphates (PIP_n_s) that are linked to p53 to regulate its stability and function (Carrillo *et al*, 2023; Chen *et al*, 2022; Choi *et al*, 2019b). Class I PITP⍺/β and the PI 4-kinase PI4KII⍺ interact with wild-type and mutant p53 in the nucleus and regulate the synthesis of p53-coupled phosphatidylinositol 4-monophosphate (PI(4)P) in response to stress (Carrillo *et al*., 2023). PIPKIα and the nuclear PI 3-kinase IPMK then bind nuclear p53 and catalyze the step-by-step generation of p53-coupled PI(4,5)P_2_ and phosphatidylinositol 3,4,5-trisphosphate (PI(3,4,5)P_3_), while the PI 3-phosphatase PTEN converts p53-PI(3,4,5)P_3_ to p53-PI(4,5)P_2_ (Chen *et al*., 2022; Choi *et al*., 2019b). Functionally, PIPn coupling to p53 recruits the small heat shock proteins (sHSPs) HSP27 and ⍺B-crystallin to the p53 complex, and these sHSPs are essential to stabilize wild-type p53 under stress and mutant p53 constitutively (Choi *et al*., 2019b). Moreover, the nuclear p53-PI(3,4,5)P_3_ complex recruits and activates PDK1, mTORC2 and Akt by a PI(3,4,5P)_3_- dependent mechanism, resulting in nuclear Akt activation, phosphorylation of the Akt substrate FOXO1/3, and inhibiting stress-induced apoptosis (Chen *et al*., 2022). These discoveries point to a nuclear PIP signaling pathway, analogous to the membrane-localized pathway, present in the membrane-free nucleus and linked to proteins such as p53.

The discovery of a nuclear PI signaling pathway coupled to nuclear proteins prompted us to postulate that a similar PI signalosome was assembled on Star-PAP to regulate its function. Consistent with this hypothesis, we report here that PITP⍺/β, PI kinases/phosphatases and sHSPs bind Star-PAP in response to stress and dynamically regulate the coupling and interconversion of PI(4)P, PI(4,5)P_2_ and PI(3,4,5)P_3_ to Star-PAP. Similar to p53 (Carrillo *et al*, 2024; Choi *et al*., 2019b), PIPn binding to Star-PAP serves as a signal to recruit sHSPs to the Star-PAP complex. KD of key components of the nuclear PI signaling pathway including PITP⍺/β, PI kinases and HSP27 inhibit protein expression levels of the Star-PAP targets HO-1 and BIK. Collectively, these data provide strong evidence for a Star-PAP-PI signalosome that regulates its activity and suggests that the nuclear PI signaling pathway is likely to encompass additional nuclear proteins beyond p53 and Star-PAP.

## Results

### Multiple PIP_n_s are coupled to Star-PAP in response to stress

Although the association of Star-PAP with PIPKI⍺ and its product PI(4,5)P_2_ was reported previously (Laishram *et al*., 2011; Li *et al*., 2012; Mellman *et al*., 2008), recent studies indicate that PIP modifying enzymes regulate the interconversion of multiple PIP_n_s that are linked to nuclear effector proteins such as p53 (Chen *et al*., 2022; Choi *et al*., 2019b). To determine if these PIP isomers are linked to Star-PAP we used specific PIP antibodies to probe Star-PAP. For this Star-PAP was ectopically expressed, was immunoprecipitated (IPed), boiled in SDS-PAPE sample buffer, separated by SDS-PAGE and then analyzed by fluorescence WB with antibodies (Abs) specific for PI(4)P, PI(4,5)P_2_, or PI(3,4,5)P_3_ (green) or HA (red) (**Fig. 1A-C**). Notably, all three of these PIP_n_s co-IPed with Star-PAP and their respective bands overlapped (yellow merged image), indicating a denaturation resistant tight linkage to Star-PAP. Microscale thermophoresis (MST) with recombinant His6-Star-PAP protein and PIPn micelles as ligands revealed high affinity interactions of PI(4)P, PI(4,5)P_2_, and PI(3,4,5)P_3_ with Star-PAP, while phosphatidylinositol did not bind Star-PAP (**Fig. EV1A**) indicating direct binding of PIPs to Star-PAP.

**Figure 1.**
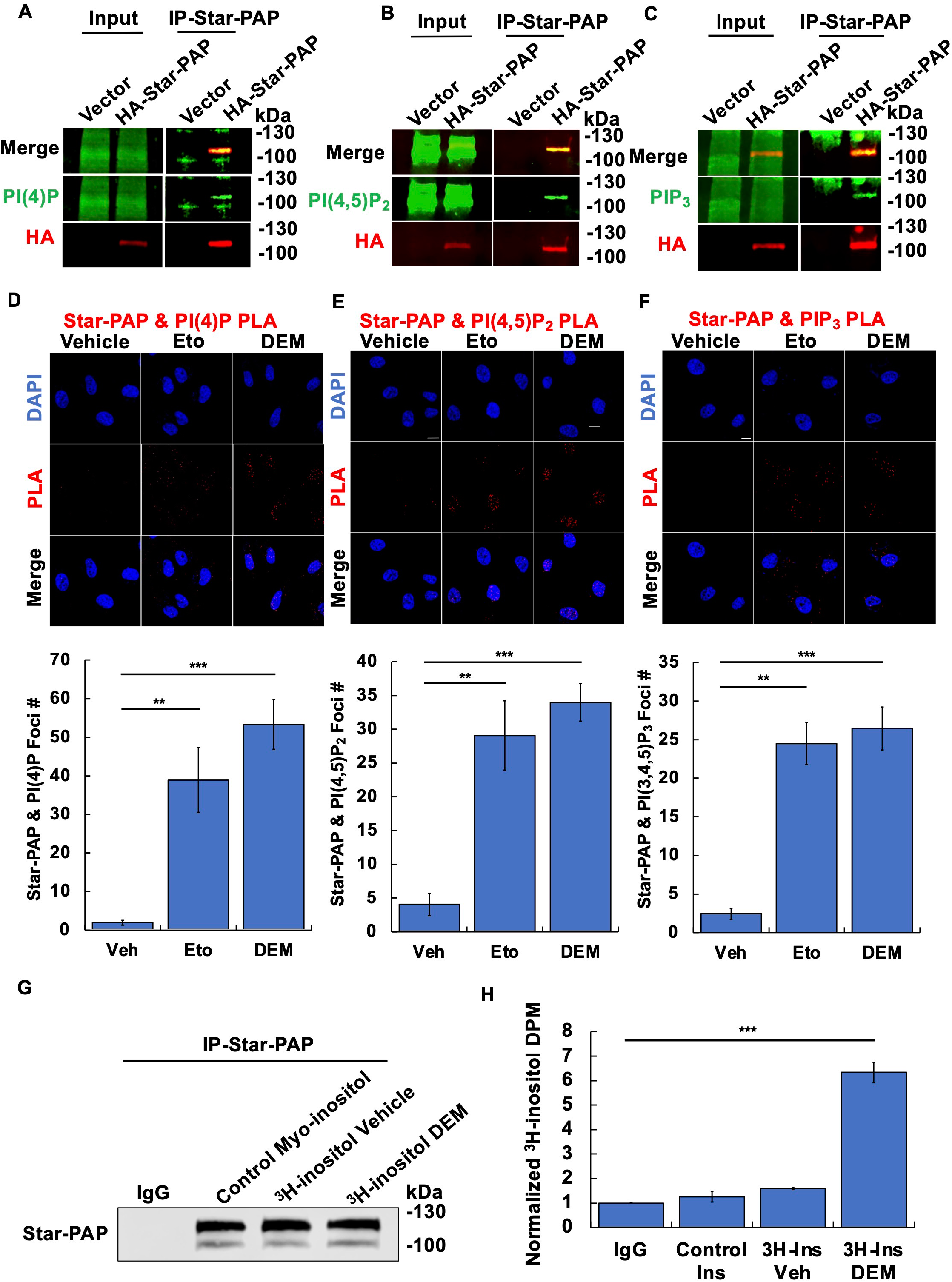
Multiple PIP_n_s are coupled to Star-PAP in response to stress. **A-C.** HEK293FT cells were transiently transfected with HA-tagged Star-PAP. 48 h later, cells were treated with 0.6% DEM for 4 h and processed for IP of Star-PAP and fluorescent WB using antibodies against HA and PI(4)P (A), PI(4,5)P_2_ (B), or PI(3,4,5)P_3_ (C). **D-F.** HeLa cells treated with 100 μM etoposide or 0.6% DEM for 4 h and PLA analysis was performed for Star-PAP and PI(4)P (C), PI(4,5)P_2_ (D), or PI(3,4,5)P_3_ (E). Images (100x with oil) were captured using a SP8 Leica confocal microscope (*n* = 3 images each from three independent experiments, Scale bar, 10 μM). PLA foci per cell in C-E were quantified using ImageJ (*n* = 30 cells from three independent experiments). *P*-values are for one-sample *t*-test versus vehicle, *** for *P* < 0.001. **G, H.** HEK293FT cells were cultured from low confluency in media containing ^3^H myo- inositol or unlabeled control myo-inositol. After 72 h of growth to confluency, cells were treated with 0.6% DEM for 4 h before being processed for IP against Star-PAP. Star-PAP level were confirmed by WB (G) before samples were resolved by SDS-PAGE and the gel lane was excised and sectioned. Gel sections were then dissolved and analyzed by liquid scintillation counting (LSC), disintegration per minute (DPM) were normalized to the IPed Star-PAP level (H). (n=3 independent experiments) *P*-values are for one-sample *t*-test versus vehicle, *** for *P* < 0.001.

To determine if Star-PAP associated with PI(4)P, PI(4,5)P_2_, and PI(3,4,5)P_3_ in HeLa cells a proximity ligation assay (PLA) was used (Chen *et al*, 2021). For these experiments, cells where fixed with formaldehyde and extensively permeabilized with detergent to remove free PIPs and then assayed for Star-PAP-PIPn complexes using Star-PAP antibodies and PIPn antibodies. The PLA data indicated that Star-PAP-PIPn complexes occur in cells, and these were greatly enhanced by genotoxic (cisplatin) and oxidative (diethyl maleate, DEM) stress (**Fig. 1D-F**). Star-PAP-PI(4,5)P_2_ complexes were largely present in the nucleus, while Star-PAP-PI(4)P/PI(3,4,5)P_3_ complexes were detected in both the nucleus and cytosol. The location of the Star-PAP-PI(3,4,5)P_3_ complexes was confirmed by expressing HA-tagged Star-PAP and using HA-antibodies for the PLA with the PI(3,4,5)P_3_ antibody (**Fig. EV1B**) and also using a distinct PI(3,4,5)P_3_ antibody (**Fig. EV1C**). These data indicate that the Star-PAP-PI(3,4,5)P_3_ complexes are both nuclear and cytosolic.

We further defined the linkage between Star-PAP and PIPs by metabolic labeling with ^3^H myo-inositol which labels specifically PIPs and inositol-phosphates in cells (Wilson & Saiardi, 2017). Both endogenous and overexpressed Star-PAP were IP’ed from metabolically labeled HEK293FT cells treated by DEM and analyzed by both WB with anti-PIP antibodies and liquid scintillation counting (LSC) to quantify ^3^H incorporation into Star-PAP (**Fig. 1G,H**, **Fig. EV1D,E**). The normalized disintegration per minute (DPM) indicated incorporation of ^3^H myo-inositol to Star-PAP under oxidative stress. As with the detection of the Star-PAP-PIPn complexes by anti-PIPn antibodies the IP’ed Star-PAP was boiled in SDS-PAPE sample buffer and separated by SDS-PAGE from free PIPs.

These findings indicate that multiple PIP_n_s are coupled to Star-PAP in response to stress.

### PI transfer proteins, PI kinases and phosphatases bind to Star-PAP in response to cell stress

Class I PITPs, PITP⍺ and PITPβ, can bind and facilitate the transfer of phosphoinositides (Hsuan & Cockcroft, 2001),have recently been shown to be required to generate the nuclear PIPn pools and initiate PIPn coupling to p53 (Carrillo *et al*., 2023; Carrillo *et al*., 2024). Further, the p53-PIPn complexes are resistant to denaturation and SDS-PAGE indicating a tight linkage like that of Star-PAP. In an ordered progression the PITPs, PI4KII⍺, PIPKI⍺, IPMK, and PTEN generate and control the interconversion of the PIP_n_s linked to p53 (Carrillo *et al*., 2023; Carrillo *et al*., 2024; Chen *et al*., 2022; Choi *et al*, 2019a). IPMK has also been shown to generate the nuclear receptor SF1 and PIP_3_ complex (Blind *et al*, 2014a; Blind *et al*, 2012). Star-PAP also associates with PITP⍺/β, PI4KII⍺, PIPKI⍺, IPMK, and PTEN in response to genomic and oxidative stress as determined by PLA (**Fig. 2A-F**). Star-PAP-PIPKI⍺ complexes were largely nuclear consistent with its role in regulating Star-PAP (Kandala *et al*, 2019; Koshre *et al*., 2021; Laishram, 2018; Mellman *et al*., 2008; Sudheesh *et al*., 2019), while each of the other Star-PAP complexes were present in the both the nucleus and cytosol. To further establish these interactions, PITP⍺/β, PI4KIIα, IPMK, and PTEN co-IPed with Star-PAP, and their interaction was enhanced by etoposide and DEM (**Fig. EV2A,B**). Ectopic expression of Myc-Star-PAP or Myc-IPMK in MDA-MB-231 cells revealed Star-PAP association with PIPKI⍺, IMPK and PTEN in response to stress largely in the nucleus (**Fig. EV2C-F**). The direct interaction of these components with Star-PAP was quantified by MST that detected high affinity interactions between Star-PAP, PITPβ and the PIP modifying enzymes (**Fig. EV2G**). Consistent with PITPβ’s role in binding and transferring phosphatidylinositol (PI), the addition of PI increased the affinity of PITPβ for Star-PAP by 10-fold (**Fig. EV2H**). These results demonstrate that multiple PITPs and PIP modifying enzymes interact with Star-PAP in a stress-regulated manner.

**Figure 2.**
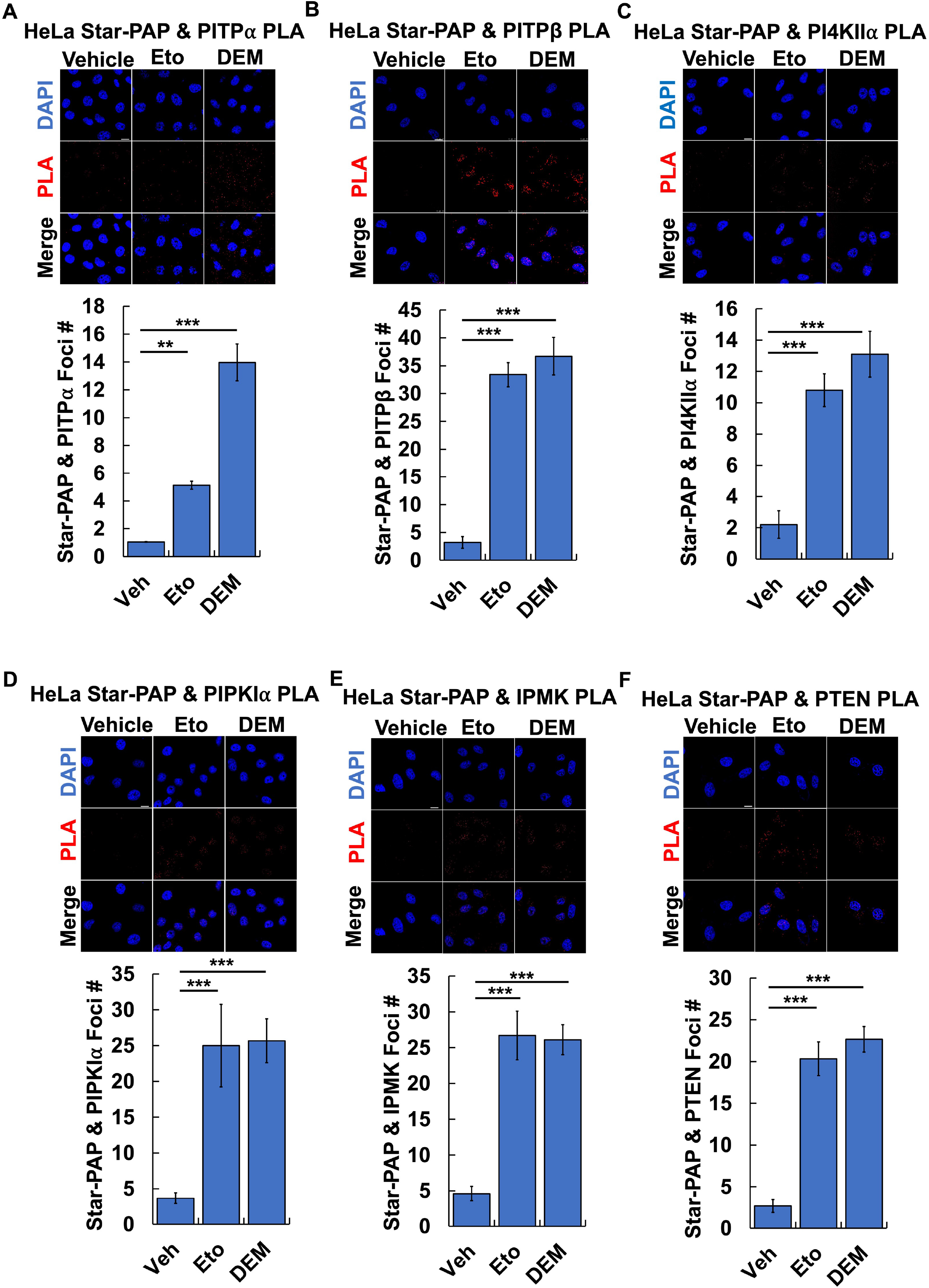
PITPs, PI kinases and phosphatases bind to Star-PAP in response to stress. **A-F.** HeLa cells treated with 100 μM etoposide or 0.6% DEM for 4 h and PLA analysis was performed for endogenous Star-PAP and class I PITPs PITP⍺ (A), PITPβ (B), PI4KII⍺ (C), PIPKI⍺ (D), IPMK (E), and PTEN (F). Images (100x with oil) were captured using a SP8 Leica confocal microscope (*n* = 3 images each from three independent experiments, Scale bar, 10 μM). PLA foci per cell were quantified using ImageJ (*n* = 30 cells from three independent experiments). *P*-values are for one-sample *t*-test versus vehicle, ** for *P* < 0.01, *** for *P* < 0.001.

### PITP⍺/β regulate PIPn and PIPKI⍺ binding to Star-PAP

To determine the role of PITP⍺/β in coupling PIP_n_s to Star-PAP, we examined the effects of combined PITP⍺/β KD on Star-PAP-PI(4)P/PI(4,5)P_2_ association in response to stress by PLA (**Fig. 3A,B**). The stress-stimulated Star-PAP-PI(4)P/PI(4,5)P_2_ complexes were virtually abolished by the combined PITP⍺/β KD. These results were confirmed by IP-WB of ectopically expressed HA-Star-PAP, which revealed markedly diminished PI4,5P_2_ co-IPed with HA-Star-PAP with either single or combined KD of PITP⍺/β (**Fig. 3C,D**). The combined KD of PITP⍺/β attenuated the interaction between PIPKI⍺ and Star-PAP as determined by co-IP under basal conditions and oxidative stress (**Fig. 3E,F**), consistent with the generation of a Star-PAP-PI(4)P complex that is then acted on by PIPKI⍺ binding to generate the STAR-PAP-PI(4,5)P_2_ complex and this aspect was explored.

**Figure 3.**
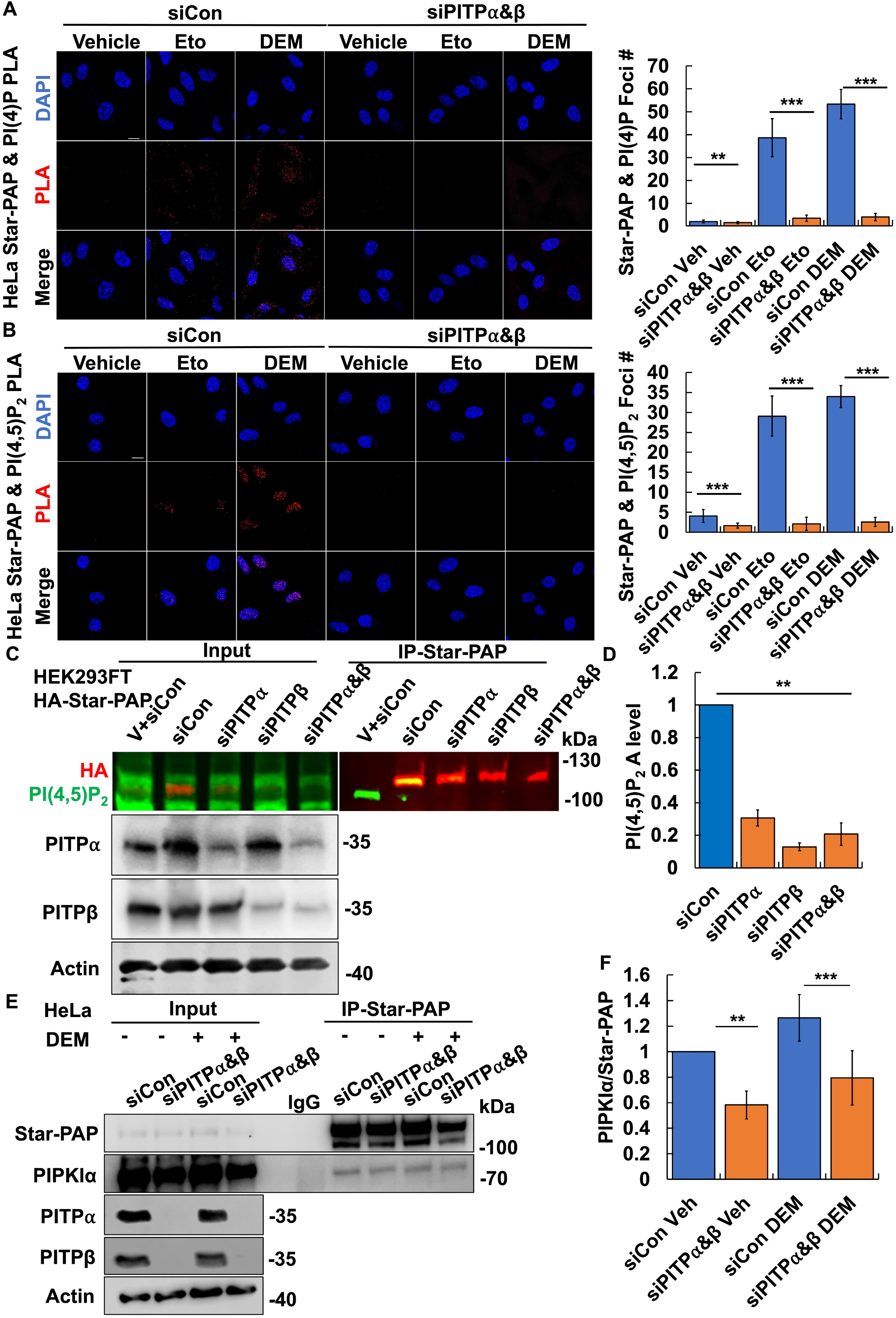
PITP⍺/β regulate PIP_n_ and PIPKI⍺ binding to Star-PAP. **A, B.** HeLa cells were transiently transfected with control siRNAs (siCon) or siRNAs targeting both PITP⍺/β. 48 h later, transfected cells were treated with 100 μM etoposide or 0.6% DEM for 4 h. PLA images were captured using SP8 Leica confocal microscope (100x with oil) for endogenous Star-PAP and PI(4)P (A) or PI(3,4,5)P_2_ (B) (*n* = 3 images each from three independent experiments, Scale bar, 10 μM). PLA foci per cell were quantified using ImageJ (*n* = 30 cells from three independent experiments). *P*-values are for one-sample *t*-test versus vehicle, ** for *P* < 0.01, *** for *P* < 0.001. **C, D.** HEK293FT cells were transiently transfected with HA-tagged Star-PAP. 48 h later, cells were transfected with control siRNAs (siCon) or siRNAs targeting PITP⍺, PITPβ or both PITP⍺/β. 48 h later, cells were processed for IP of Star-PAP and fluorescent WB for using antibodies against HA or PI(4,5)P_2_ (C) (*n* = 3 independent experiments). PI(4,5)P_2_ levels were quantified by Odyssey Imaging System (LI-COR Biosciences) and normalized to the HA level in each condition (D). *P*-values are for one-sample *t*-test versus control siRNA, ** for *P* < 0.01. **E, F.** HeLa cells were transfected with control siRNAs or siRNAs targeting both PITP⍺/β. 48 h later, cells were treated with 0.6% DEM for 4 h and processed for IP of Star-PAP and WB using antibodies against Star-PAP or PIPKI⍺ (E) (*n* = 3 independent experiments). PIPKI⍺ levels were quantified by Odyssey Imaging System (LI-COR Biosciences) and normalized to the Star-PAP level in each condition (F). *P*-values are for one-sample *t*-test versus control siRNA, ** for *P* < 0.01, *** for *P* < 0.001.

### PI4KII⍺, PIPKI⍺, IPMK and PTEN regulate the interconversion of Star-PAP- PIP_n_ complexes

The KD of the PITPs results in a loss of the Star-PAP-PI(4)P and PI(4,5)P_2_ complexes. To examine whether Star-PAP-PI4P is generated by PI4KII⍺ and then converted to Star-PAP-PI4,5P_2_ by PIPKI⍺, these PIPKs were KD’ed and the Star-PAP-PIPn complexes quantified by PLA (**Fig. 4A-D**, **Fig. EV4A-D**). Consistent with the interaction with Star-PAP, the PI4KII⍺ KD reduced both the Star-PAP-PI4P and Star-PAP-PI4,5P_2_ complexes. Consistently, PIPKI⍺ KD increased Star-PAP-PI4P complexes but diminished the Star-PAP-PI4,5P_2_ complexes. As both IPMK and PTEN bind to Star-PAP, we next determined the roles of the PI3K IMPK and the PI(3,4,5)P_3_ 3-phosphatase PTEN in regulating Star-PAP-PI(3,4,5)P_3_ complexes. The KD of PIPKI⍺ or PTEN reduced Star-PAP-PI4,5P_2_ complexes determined by PLA, while IMPK KD increased Star-PAP-PI(4,5)P_2_ levels (**Fig. 4E**). Moreover, KD of PIPKI⍺ or IMPK reduced Star-PAP-PI(3,4,5)P_3_ foci, while PTEN KD increased Star-PAP-PI(3,4,5)P_3_ complexes (**Fig. 4F**). KD of PIPKI⍺ KD and IPMK in a second cancer line confirmed the observed reduction of Star-PAP-PI(3,4,5)P_3_ PLA foci (**Fig. EV4E,F**). These data are consistent with recent studies have shown that IPMK and PTEN regulate the interconversion of PI(4,5)P_2_ and PI(3,4,5)P_3_ coupled to SF1 and p53 (Blind *et al*, 2014b; Chen *et al*., 2022). Combined, these data indicate that Star-PAP- PI(4)P is synthesized by PI4KII⍺, requiring the PITPs, and is then sequentially phosphorylated to Star-PAP-PI(4,5)P_2_ and Star-PAP-PI(3,4,5)P_3_ by PIPKI⍺ and IPMK, respectively. The Star-PAP-PI(3,4,5)P_3_ complex is dephosphorylated to Star-PAP- PI(4,5)P_2_ by PTEN.

**Figure 4.**
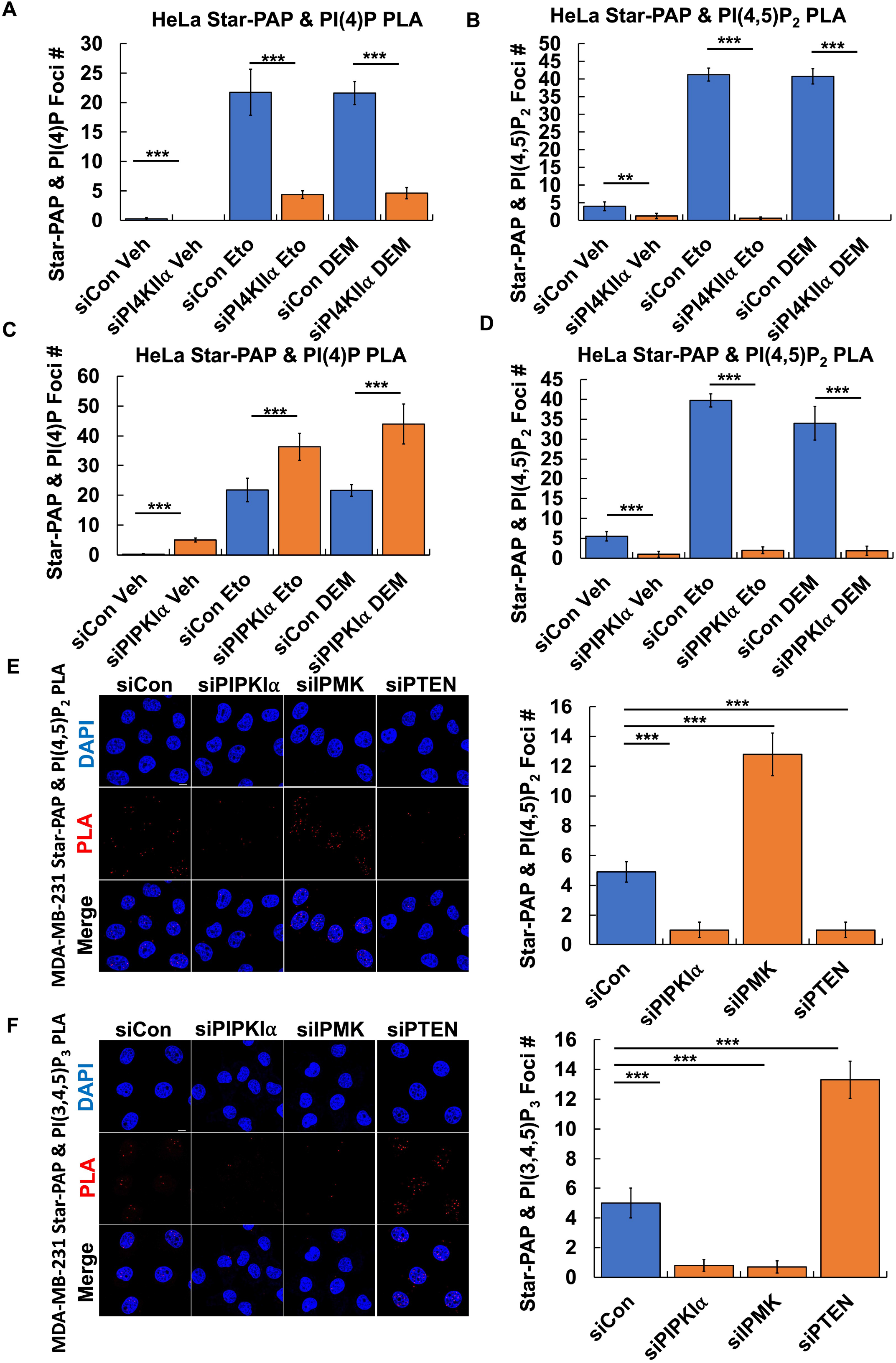
PI4KII⍺, PIPKI⍺, IPMK, and PTEN regulate the interconversion of Star-PAP- coupled PIP_n_s. **A, B.** HeLa cells were transfected with control siRNAs or siRNAs targeting PI4KII⍺. 48 h later, cells were treated with 100 μM etoposide or 0.6% DEM for 4 h before performing PLA analysis of Star-PAP and PI(4)P or PI(4,5)P_2_ (*n* = 3 images each from three independent experiments, Scale bar, 10 μM). PLA foci of Star-PAP-PI(4)P (A) and Star- PAP-PI(4,5)P_2_ (B) per cell were quantified using ImageJ (*n* = 30 cells from three independent experiments). *P*-values are for one-sample *t*-test versus vehicle, ** for *P* < 0.01, *** for *P* < 0.001. **C, D.** HeLa cells were transfected with control siRNAs or siRNAs targeting PIPKI⍺. 48 h later, cells were treated with 100 μM etoposide or 0.6% DEM for 4 h before performing PLA analysis of Star-PAP and PI(4)P or PI(4,5)P_2_ (*n* = 3 images each from three independent experiments, Scale bar, 10 μM). PLA foci of Star-PAP-PI(4)P (C) and Star- PAP-PI(4,5)P_2_ (D) per cell were quantified using ImageJ (*n* = 30 cells from three independent experiments). *P*-values are for one-sample *t*-test versus vehicle, *** for *P* < 0.001. **E, F.** MDA-MB-231 cells were transfected with control siRNAs or siRNAs targeting PIPKI⍺, IPMK or PTEN. 48 h later, PLA analysis was performed on Star-PAP and PI(4,5)P_2_ (E) or PI(3,4,5)P_3_ (F) (*n* = 3 images each from three independent experiments, Scale bar, 10 μM). PLA foci per cell were quantified using ImageJ (*n* = 10 cells from three independent experiments). *P*-values are for one-sample *t*-test versus control siRNA, *** for *P* < 0.001.

### Small heat shock proteins bind Star-PAP by a PIP_n_-regulated mechanism

The small heat shock proteins (sHSPs) HSP27 and ⍺B-crystallin bind p53 and their binding is controlled PI(4,5)P_2_ that bind to the p53 C-terminal polybasic sequence (Choi *et al*., 2019b). We examined whether these sHSPs are regulated similarly in response to the PIP_n_ linkage to Star-PAP. Consistently, HSP27 and ⍺B-crystallin associated with Star-PAP in HeLa cells stimulated with etoposide or DEM as determined by PLA (**Fig. 5A,B**). Stress also stimulated the interaction between Star-PAP and these sHSPs as determined by co-IP (**Fig. 5C**) and PIPKI⍺ KD disrupted HSP27 binding to Star-PAP in response to stress (**Fig. 5D**). MST and *in vitro* binding assays between His-tagged Star-PAP and HSP27/⍺B-crystallin purified from *E. coli* indicated direct interaction between Star-PAP and the small heat shock proteins (**Fig. EV5A-D**). Notably, the addition of PI4P, PI4,5P_2_ or PI3,4,5P_3_ micelles, but not phosphatidylinositol, enhanced the interaction of HSP27 with Star-PAP; PI(4,5)P_2_ and PI(3,4,5)P_3_ increased the binding affinity ∼10 fold (**Fig. EV5A,B**). Moreover, HSP27 did not bind directly to PI4P or PI4,5,P_2_ and bound only modestly to PI3,4,5P_3_, indicating that PIPn binding to Star-PAP is regulating the Star-PAP- HSP27 interaction (**Fig. EV5B**). The effect of PI(4,5)P_2_ on Star-PAP-HSP27 binding was further confirmed by *in vitro* binding assay in which a constant amount of His-tagged Star-PAP and HSP27 were mixed with an increasing concentration of PI(4,5)P_2_ before performing IP of Star-PAP. Addition of 2 μM PI4,5P2 increased HSP27 binding to Star-PAP more than 10-fold (**Fig. EV5E**). PIPKI⍺ KD (**Fig. 5D**) and combined PITP⍺/β KD inhibited the Star-PAP-HSP27 interaction (**Fig. EV5F**), further implicating PIPn coupling to Star-PAP as the signal that recruits sHSPs to the complex.

**Figure 5.**
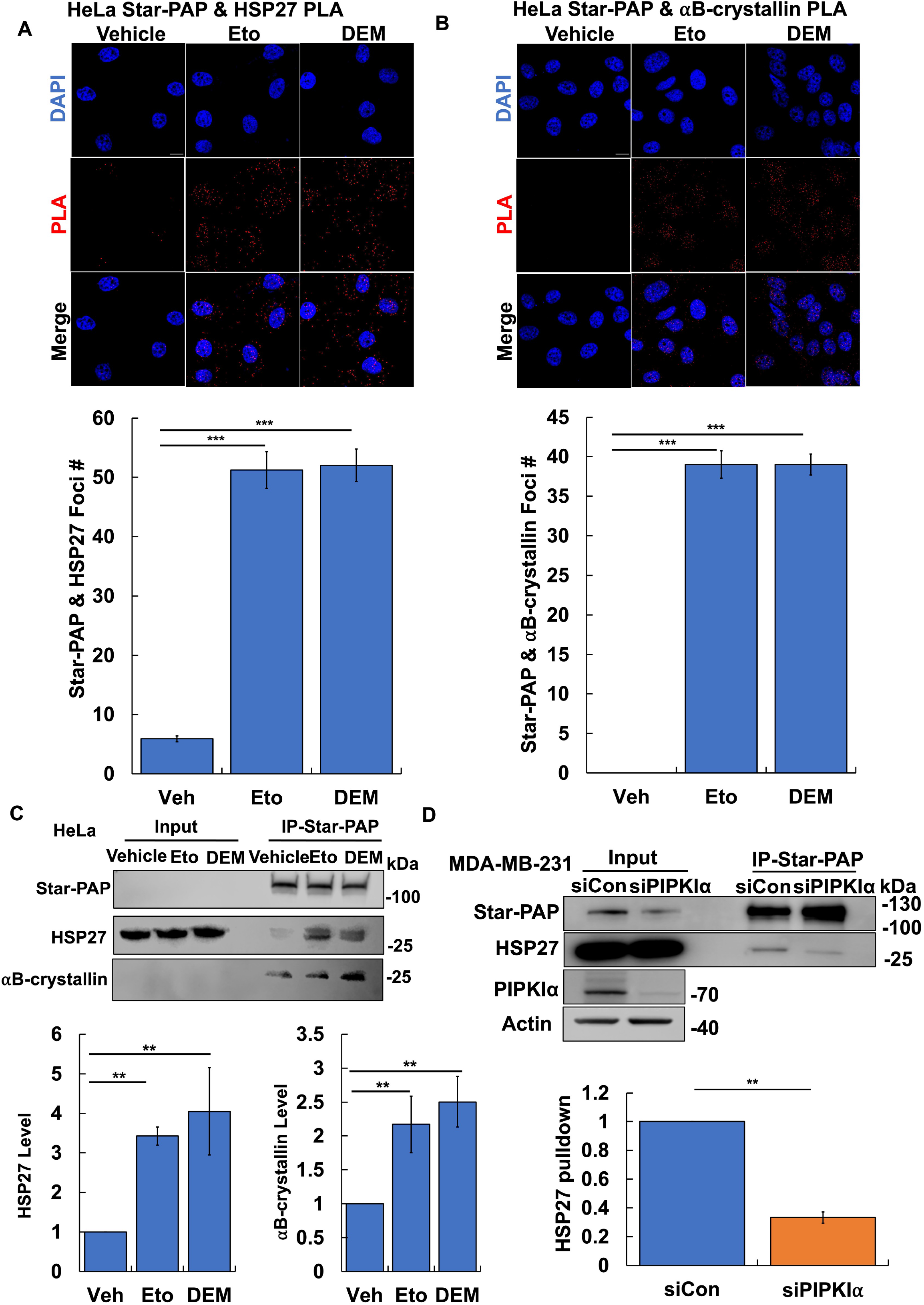
Small heat shock proteins bind Star-PAP in response to stress. **A, B.** HeLa cells treated with 100 μM etoposide or 0.6% DEM for 4 h. 48 h later, PLA analysis was performed for Star-PAP and HSP27 (A) or ⍺B-crystallin (B) (*n* = 3 images each from three independent experiments, Scale bar, 10 μM). PLA foci per cell were quantified using ImageJ (*n* = 30 cells from three independent experiments). *P*-values are for one-sample *t*-test versus vehicle, *** for *P* < 0.001. **C.** HeLa cells were treated with 100 μM etoposide or 0.6% DEM for 4 h and processed for IP of Star-PAP and WB using antibodies against Star-PAP, HSP27, and ⍺B-crystallin (*n* = 3 independent experiments). HSP27 and ⍺B-crystallin levels were quantified by Odyssey Imaging System (LI-COR Biosciences) and normalized to the Star-PAP level in each condition. *P*-values are for one-sample *t*-test versus vehicle, ** for *P* < 0.01. **D.** HeLa cells were transfected control siRNAd or siRNAs targeting PIPKI⍺. 48 h later, cells were processed for IP of Star-PAP and WB with antibodies against Star-PAP and HSP27 (*n* = 3 independent experiments). HSP27 levels were quantified and normalized to the Star-PAP level in each condition. *P*-values are for one-sample *t*-test versus control siRNA, ** for *P* < 0.01.

### PITP⍺/β, PI4KIIα, HSP27, and IPMK regulate expression of Star-PAP targets in response to stress

Star-PAP polyadenylation targets downstream of genotoxic or oxidative stress are regulated by PIPKI⍺ and PI(4,5)P_2_ recruitment to Star-PAP (Laishram *et al*., 2011; Li *et al*., 2012; Mellman *et al*., 2008; Mohan *et al*, 2015). We investigated whether more recently discovered pathway components (PITP⍺/β, PI4KIIα, HSP27 and IPMK) regulated the Star-PAP targets HO-1 and BIK. Combined PITP⍺/β KD in HeLa and HEK293FT cells reduced DEM-stimulated HO-1 protein expression, like that of Star-PAP KD (**Fig. 6A,B**, **Fig. EV6A,B**). These KD’s also reduced basal HO-1 levels. Moreover, KD of PI4KIIα, HSP27, IPMK or Star-PAP all reduced HO-1 protein levels in response to oxidative stress and BIK levels in response to genotoxic stress (**Fig. 6C-J**, **Fig. EV6C-J**). These data demonstrate that the PIPn coupling to Star-PAP and the requisite enzymatic/chaperone machinery of the stress-regulated polyadenylation process of target mRNAs that are dependent on Star-PAP for their expression.

**Figure 6.**
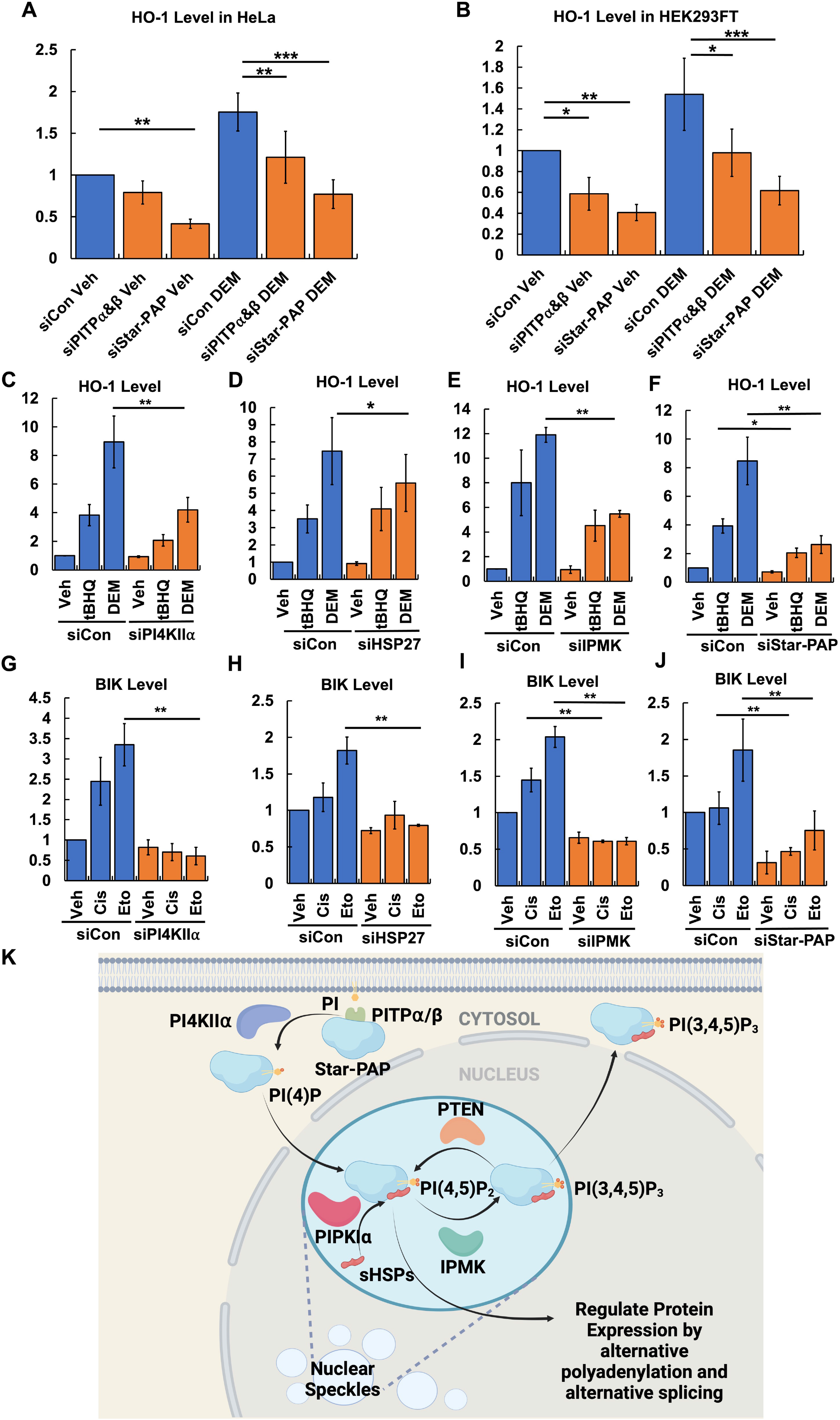
PITP⍺/β, HSP27, and IPMK regulate expression of Star-PAP targets in response to stress. **A.** HeLa cells were transfected with control siRNAs or siRNAs targeting both PITP⍺/β or Star-PAP. 48 h later, cells were treated with 0.6% DEM for 4 h and then processed for WB using an antibody against HO-1 (*n* = 3 independent experiments). HO-1 levels were quantified by Odyssey Imaging System (LI-COR Biosciences) and normalized to the actin level in each condition. *P*-values are for one-sample *t*-test versus control siRNA, ** for *P* < 0.01, *** for *P* < 0.001. **B.** HEK293FT cells were transfected with control siRNAs or siRNAs targeting both PITP⍺/β or Star-PAP. 48 h later, cells were treated with 0.6% DEM for 4 h and then processed for WB with an antibody against HO-1 (*n* = 3 independent experiments). HO- 1 levels were quantified and normalized to the actin level in each condition. *P*-values are for one-sample *t*-test versus control siRNA, * for *P* < 0.05, ** for *P* < 0.01, *** for *P* < 0.001. **C-F.** MDA-MB-231 cells were transfected with control siRNAs or siRNAs targeting PI4KII⍺ knockdown (C), HSP27 knockdown (D), IPMK knockdown (E), or Star-PAP knockdown (F). 48 h later, cells were treated with 1 mM tBHQ or 0.6% DEM for 4 h and then processed for WB with an antibody against HO-1 (*n* = 3 independent experiments). HO-1 levels were quantified and normalized to the actin level in each condition. *P*-values are for one-sample *t*-test versus control siRNA, * for *P* < 0.05, ** for *P* < 0.01. **G-J.** MDA-MB-231 cells were transfected with control siRNAs or siRNAs PI4KII⍺ knockdown (G), HSP27 knockdown (H), IPMK knockdown (I), or Star-PAP knockdown (J). 24 h later, cells were treated with 30 μM Cisplatin for 24 h or 100 μM etoposide for 4 h and then processed for WB using an antibody against BIK (*n* = 3 independent experiments). BIK levels were quantified and normalized to the actin level in each condition. *P*-values are for one-sample *t*-test versus control siRNA, * for *P* < 0.05, ** for *P* < 0.01. **I.** Model depicting the nuclear PI signalosome on Star-PAP. Created by Biorender.

## Discussion

Star-PAP has been previously been reported to bind PIPKI⍺ and PI(4,5)P_2_ and this regulates it’s poly(A) polymerase activity (Mellman *et al*., 2008). Here, we demonstrate that Star-PAP is tightly linked to the PI(4)P, PI(4,5)P_2_ and PI(3,4,5)P_3_ isomers, and this linkage is resistant to denaturation and SDS-PAGE (**Fig. 1A-H**) and are assembled on Star-PAP in response to stress (**Fig. 6K**). The Star-PAP-PIPn complexes are initiated by the class I PI transfer proteins, PITP⍺/β, which transport phosphatidylinositol from cytosolic membranes to the nucleus in response to stress (Carrillo *et al*., 2023; Carrillo *et al*., 2024). Indeed, the localization of Star-PAP-PITP⍺/β and Star-PAP-PI(4)P complexes in both the cytosol and nucleus suggests the origin of the PIPn coupled to Star-PAP is from cytoplasmic membranes, while PIPKI⍺ binding, PI(4,5)P_2_ coupling, and sHSP recruitment are largely in the nucleus. Notably, PI increases the binding affinity of PITPs for Star-PAP by 10-fold, underscoring the critical role of PITP binding to PI in regulating its interaction with Star-PAP. The combined KD of PITP⍺/β virtually eliminates the stress-induced coupling of PI(4)P and PI(4,5)P_2_ to Star-PAP as well as its interaction with PIPKI⍺, underscoring the essential roles of PITP⍺/β in mediating these events and indicates redundant functions,. The PI kinases PI4KII⍺, PIPKI⍺ and IMPK and the 3- phosphatase PTEN each bind Star-PAP with high affinity and regulate the interconversion of PIP_n_s linked to Star-PAP in response to stress. These coupled PIP_n_s enhance the interaction of Star-PAP with the sHSPs, both HSP27 and ⍺B-crystallin, which like the PITPs and PI kinases are required for Star-PAP-dependent regulation of its targets HO-1 and BIK. Overall, our data point to a Star-PAP-PI signalosome that controls PI coupling and Star-PAP-dependent regulation of its mRNA targets in response to stress.

The nuclear Star-PAP-PI signalosome reported here resembles the analogous pathway recently described for p53 (Carrillo *et al*., 2023; Carrillo *et al*., 2024; Chen *et al*., 2022; Choi *et al*., 2019b). Here, the pathway is initiated by PITP⍺/β binding to its target, which is dramatically enhanced by phosphatidylinositol loading of the PITPs (**Fig. 2G,H**). The PIP kinases, PTEN phosphatase and sHSPs identified here also play a role in regulating the interconversion of linked PIP_n_s on p53 (Chen *et al*., 2022; Choi *et al*., 2019b). However, unlike p53 where the generation of p53-PI(3,4,5)P_3_ has been linked to nuclear Akt activation (Chen *et al*., 2022). The Star-PAP-PI(3,4,5)P_3_ complex is different in that it is both nuclear and cytoplasmic suggesting that Star-PAP-PI(3,4,5)P_3_ has a role in both compartments (**Fig. EV4E,F**) but this maybe cell type or signal regulated as the Star-PAP- PI(3,4,5)P_3_ complex in MDA-MB-231 is predominantly nuclear (**Fig. 4F**). Yet, it is clear that the interconversion of Star-PAP-PIPn complexes is regulated by PIP kinases and phosphatases in an ordered fashion and this in turn is regulated by cell stress (**Fig. 6K**).

The data presented here support that PIP_n_s are tightly linked to Star-PAP, namely, the ability to WB PI(4)P, PI(4,5)P_2_ or PI(3,4,5)P_3_ following Star-PAP IP by a process that requires boiling the samples in SDS buffer. Compelling evidence comes from the ability to detect incorporated ^3^H-inositol label in Star-PAP after metabolic labeling of cells with ^3^H-myo-inositol. These findings are consistent with a covalent linkage indicating a post-translational modification (PTM). This indicates that Star-PAP is regulated by a linkage of PIPn isomers likely starting with PI from the PITP:PI complexes. This is a distinct mechanism for PIPn control of proteins. For example, the nuclear receptor steroidogenic factor 1 (SF-1) binds PI(4,5)P_2_ and PI(3,4,5)P_3_ and these bound PIP_n_s are interconverted by IPMK and PTEN in an analogous fashion (Blind *et al*., 2012; Bryant & Blind, 2019). However, in this case the PIP_n_s are bound to SF-1 in a mechanism like that of PI binding to the PITPs, through a hydrophobic pocket where the PIP acyl chains are buried and the PIPn head group is exposed and can be acted on by PIP enzymes. In the Star-PAP-PIPn linkage model the PITPα/β, PI4KIIα, PIPKIα, IPMK and PTEN bind to E. coli expressed Star-PAP that lack the PIP linkage. This is a key difference as these specific interactions define the signaling specificity for the Star-PAP-PIPn complexes and these complexes then have functions that are defined by the specific PIPn that is linked to Star-PAP. This general mechanism could be expanded so that all seven of the PIPn isomers could be specifically generated by protein-protein interactions that recruit specific PIP kinases and phosphatases. The action of phospholipase C, known to be nuclear (Barlow *et al*, 2012; Cocco *et al*, 2009), on the PIPn complexes could be another mechanism to generate additional messages.

Our data point to a fundamentally new paradigm of nuclear PI signaling entirely distinct from their canonical role in membranes as lipid second messengers. In the nuclear PI pathway, PIP_n_s are linked to nuclear proteins as a putative PTM that requires the nuclear translocation of PITP⍺/β to initiate the process. A nuclear PI signalosome or scaffolding complex is assembled by the target protein that enables exquisite spatial and temporal synthesis of multiple distinct PIPn isomers, each with potentially distinct binding partners and effector pathways as exemplified by the p53-PI(3,4,5)P_3_ complex, which recruits and activates nuclear Akt (Chen *et al*., 2022). It is also interesting to notice that IPMK depletion has been reported to trigger nuclear accumulation of Poly(A)^+^ RNA and decrease the nuclear export of a subset of mRNA transcripts (Wickramasinghe *et al*, 2013). These findings suggested that the Star-PAP-PI(3,4,5)P_3_-IPMK complex may be involved in the nuclear export pathways downstream of the polyadenylation process. Consistent with this concept is evidence indicating that Akt can regulate mRNA export (Quaresma *et al*, 2013). As it is found that in hypertrophic cardiac cells and breast cancer cells, Star-PAP expression is suppressed (Sudheesh *et al*., 2019; Yu *et al*, 2017), this phosphoinositide signalosome pathway mediating Star-PAP targets provides new potential therapeutic targets in a variety of disease.

## Materials and Methods

### Cells

HeLa, HEK293FT, and MDA-MB-231 cells were purchased from ATCC. Cells were maintained in DMEM (#10-013-CV, Corning) supplemented with 10% fetal bovine serum (#SH30910.03, Hyclone) and 1% penicillin/streptomycin (#15140-122, Gibco). The cell lines used in this study were routinely tested for mycoplasma contamination using MycoAlert Mycoplasma Detection Kit (#LT07-318, Lonza), and mycoplasma-negative cells were used. None of the cell lines used in this study are listed in the database of commonly misidentified cell lines maintained by ICLAC. The Star-PAP HA-tagged construct used for this work is described previously (Mohan *et al*., 2015). Constructs were transfected into HEK293FT cells by branched polyethylenimine (bPEI, #408727, Sigma-Aldrich). 15 μg of DNA and 20 μl of bPEI were mixed in 1 ml of PBS for each 10-cc dish and incubated in 37°C for 20 min before adding to the culture medium. HeLa cells were transfected lipid-based delivery system from Invitrogen (Lipofectamine™3000, #L300015, Thermo Fisher Scientific) according to the manufacturer’s instructions. 3 μg of DNA and 6 μl of lipid were used for each well of cells in 6-well plates.

### Antibodies and Reagents

Monoclonal antibodies against HA-tag (clone C29F4, #3724, Cell Signaling) HSP27 (clone D6W5V, #95357, Cell Signaling; clone G31, #2402, Cell Signaling), αB-crystallin (clone D6S9E, #45844, Cell Signaling), HO-1 (clone D60G11, #5853, Cell Signaling), and polyclonal antibodies against β-actin (#4967, Cell Signaling), PITPα (#16613-1-AP, ThermoFisher), PITPβ (#ab127563, abcam), PIPKIα (#9693, Cell Signaling) BIK (#4592, Cell Signaling) were used in this study. Polyclonal antibodies against Star-PAP, PIPKIα, and IPMK were produced as described previously (Mellman *et al*., 2008; Choi *et al*., 2016). For conventional immunostaining and PLA analysis of phosphoinositides, anti- PI4P (#Z-P004), PI4,5P2 (#Z-P045) and PI3,4,5P_3_ (#Z-P_3_45) antibodies were purchased from Echelon Biosciences. For Western blots (WB), antibodies were diluted at a 1:1000 ratio except for Star-PAP (1:200). For immunoprecipitation (IP), nProtein A Sepharose™ 4 Fast Flow resin (#17528001, Cytiva) was used together with Star-PAP antibody. For immunostaining analyses and proximity ligation assay (PLA), antibodies were diluted at a 1:100 ratio. For the knockdown (KD) experiments, custom siRNA targeting the 3’UTR of Star-PAP (sense 5’-GUGUGUUUGUCAGUGGCUUdTdT-3’ and anti-sense 5’- AAGCCACUGACAAACACACdTdT-3’), the ON-TARGETplus siRNA SMARTpool with 4 siRNAs in combination against human PITPα (#L-018010-00), PITPβ (#L-006459-01), HSP27 (#L-005269-00), PIPKIα (#L-041183-00), IPMK (#L-006740-00) were purchased from Dharmacon. Non-targeting siRNAs (Dharmacon) were used as a control. The siRNAs were delivered to HEK293FT cells by bPEI or HeLa and MDA-MB-231 cells by RNAiMAX reagent (#13778150, Thermo Fisher Scientific) KD efficiency was determined by WB. KD efficiency greater than 80% was required to observe phenotypic changes in the study. Cisplatin (Cis, #NC1706394, Fisher Scientific), etoposide (eto, #S1225, Sellekchem), tert-Butylhydroquinone (tBHQ, #SPCM-T1540-06, Spectrum), and diethyl maleate (DEM, #D97703-500G, Sigma-Aldrich) were used as cellular stressors.

### Immunoprecipitation (IP) and Western blotting (WB)

Cells were removed from the medium after the indicated treatment, washed three times with ice-cold PBS, and lysed in an ice-cold RIPA lysis buffer system (#sc-24948, Santa Cruz Biotechnology) with 1x Pierce Protease and Phosphatase Inhibitor Mini Tablet (#A32959, Thermo Fisher Scientifics). The cell lysates were then sonicated at 15% amplitude for 15 s. After sonication, the cell lysates were centrifuged at maximum speed for 10 min to collect the supernatant. The protein concentration in the supernatant was measured by the Bradford protein assay (#5000201, BIO-RAD) according to the manufacturer’s instructions. Equal amounts of protein were used for further analysis. All antibodies were diluted at a 1:1000 ratio for WB unless otherwise indicated. For IP, 0.5- 1 mg of cell lysate was incubated with 4 μg of Star-PAP antibody 4°C for 1-4 h and add 20 μl of nProtein A Sepharose, incubating in 4°C overnight. After washing three times with PBST (PBS with 0.1% Tween 20), the protein complex was eluted with SDS sample buffer. The sample was then boiled at 95°C for 5 min. For WB, 5-20 μg of protein were loaded. For WB of IP complexes, IgG control and mouse primary antibodies were used to avoid non-specific detection of immunoglobulin in the IPed samples. WBs were developed and quantified by Odyssey Imaging System (LI-COR Biosciences). The unsaturated exposure of WB images was used for quantification with the appropriate loading controls as standards. Statistical data analysis was performed with Microsoft Excel, using data from at least three independent experiments.

### Fluorescent IP-WB

HEK293FT Cells were transfected with HA-Star-PAP then lysed in a RIPA lysis buffer system after the indicated treatment and quantified for protein concentration as described above. For HA-tagged Star-PAP IP, 0.5-1 mg of cell lysate was incubated with 4 μg of Star-PAP antibody 4°C for 1-4 h and add 20 μl of nProtein A Sepharose, incubating in 4°C overnight. Mixture of Star-PAP antibody and Sepharose in cell lystate transfected with empty pCMV-HA vectors was used as control IgG. After washing three times with PBST, the protein complex was eluted with SDS sample buffer. The sample was then boiled at 95°C for 5 min. For WB, 5-20 μg of protein were loaded. The protein complexes associated with Star-PAP were resolved by SDS-PAGE and transferred onto a PVDF membrane (#IPVH00010, MilliporeSigma). The membrane was blocked with 1% BSA in TBS for 1 h at room temperature. For double fluorescent IP-WB detecting HA-Star-PAP- PI4P/PI4,5P_2_/PI3,4,5P_3_ complexes, anti-HA rabbit monoclonal IgG antibody at 1:2000 dilution and anti-PI4P mouse monoclonal IgM antibody, PI4,5P_2_ mouse monoclonal IgM antibody, or PI3,4,5P_3_ mouse monoclonal IgM antibody at 1:1000 dilution were mixed together in blocking buffer with 0.02 % Sodium Azide and incubate with the membrane at 4°C overnight. The next day, the membrane was washed three times with TBST for 10 min each time. For the secondary antibody incubation, goat anti-rabbit IgG antibody conjugated with IRDye 800CW fluorophore (#926-32211, LI-COR) detectable on the 800 nm wavelength channel of the Odyssey Fc Imaging System (LI-COR Biosciences) and goat anti-mouse IgM antibody conjugated with IRDye 680RD fluorophore (#926-68180, LI-COR) detectable on the 700 nm wavelength channel at 1:10000 dilution were mixed in blocking buffer with 0.01 % SDS and 0.1% Tween 20 and incubated with the membrane at room temperature for 2 h. The membrane was then washed three times with TBST for 10 min each time. The images were subsequently acquired simultaneously using the 700 and 800 nm wavelength channels on the Odyssey Fc Imaging System (LI-COR Biosciences). The HA-Star-PAP-associated PI4P/PI4,5P_2_/PI3,4,5P_3_ complex was visualized by overlapping the 700 and 800 nm channels. Statistical data analysis was performed with Microsoft Excel, using data from at least three independent experiments.

### Immunofluorescent (IF) and Confocal Microscopy

For immunofluorescence studies, cells were grown on coverslips coated with 0.2 % gelatin (#G9391, MilliporeSigma). Cells were fixed with 4% paraformaldehyde (PFA) (#sc- 281692, Santa Cruz Biotechnology) for 20 min at room temperature followed by washing three times with PBS. Next, the cells were permeabilized with 0.3% Triton-X100 for 30 min to thoroughly permeate the nuclear envelope and rewashed three times with PBS. The cells were then blocked with 1% BSA in PBS for one hour at room temperature. After blocking, cells were incubated with primary antibodies overnight at 4°C. The cells were then washed three times with PBS and incubated with Goat anti-Mouse IgG (H+L) Cross Absorbed Secondary Antibody, Alexa Fluor™ 555 (#A-21422, Thermo Fisher Scientifics) and Goat anti-Rabbit IgG (H+L) Cross Absorbed Secondary Antibody, Alexa Fluor™ 488 (#A-11008, Thermo Fisher Scientifics) for 1 hour at room temperature. After secondary antibody incubation, the cells were washed three times. Then the coverslips were taken out from the dishes, air-dried for 10 min at room temperature and mounted in ProlongTM Glass Antifade Mountant with NucBlueTM Stain (#P_3_6985, Thermo Fisher Scientific). The images were taken by Leica SP8 3xSTED Super-Resolution Microscope, which is both a point scanning confocal and 3xSTED super-resolution microscope. The Leica SP8 3xSTED microscope was controlled by LASX software (Leica Microsystems). All images were acquired using the 100X objective lens (N.A. 1.4 oil). The z-stack images were taken with each frame over a 0.2 μm thickness. For quantification, the mean fluorescent intensity of channels in each cell was measured by LASX. GraphPad Prism Version 9.4.1 generated quantitative graphs. The images were processed using ImageJ.

### Proximity Ligation Assay (PLA)

PLA was utilized to detect *in situ* protein-protein/PI interaction as previously described (Chen *et al*., 2022; Chen *et al*., 2021; Choi *et al*., 2019). After fixation and permeabilization, cells were blocked before incubation with primary antibodies as in routine IF staining. The cells were then processed for PLA (#DUO92101, MilliporeSigma) according to the manufacturer’s instruction and previous publications (Chen *et al*., 2022; Chen *et al*., 2021; Choi *et al*., 2019). Post-PLA slides were further processed for immunofluorescent staining against the nuclear membrane marker (Lamin A/C, #8617, Cell Signaling). After washing three times with PBST, the slides were mounted with ProlongTM Glass Antifade Mountant with NucBlueTM Stain (#P_3_6985, Thermo Fisher Scientific). The Leica SP8 confocal microscope detected PLA signals as discrete punctate foci and provided the intracellular localization of the complex. ImageJ was used to quantify the nuclear PLA foci. Statistical data analysis was performed with Microsoft Excel, using data from at least three independent experiments.

### *In vitro* Binding Assay

The recombinant proteins Star-PAP, HSP27, and αB-crystallin used for *in vitro* binding assays were purified from BL21 *E*. coli cells. The plasmids in bacterial-expression backbone were transfected into BL21 cell and grown in LB at 37°C overnight, followed by 0.5μM Isopropyl β-D-1-thiogalactopyranoside (IPTG) induction at room temperature overnight. The cells were lysed by 1% Brij58 lysis buffer (150 mM NaCl, 20 mM Tris, 1% Brij58, 2mM MgCl2, 2mM CaCl2, pH 7.4 ∼ 7.5, 1 tablet of cOmplete Mini, EDTA-free protease inhibitor cocktail per 10 ml of buffer, 10 μg/ml DNase, 20 μg/ml RNase, 50 μg/ml Lysozyme) and incubated with Ni-NTA Agarose () 4°C overnight. After washing three times with washing buffer (10 mM Imidazole, 300 mM NaCl, 50 mM Tris-Cl, 0.5% Triton X-300), the proteins were eluted from the column with elution buffer (250 mM Imidazole, 300 mM NaCl, 50 mM Tris-Cl, 0.5% Triton X-300). The eluted proteins were dialyzed in PBS using Slide-A-Lyzer© Dialysis Cassette, 10.000 MWCO (). The binding assay was performed in PBS by incubating a constant amount of Star-PAP with an increasing amount of HSP27/αB-crystallin in the presence of 4 μg of Star-PAP antibody and 20 μl of nProtein A Sepharose. After incubating overnight at 4°C, unbound proteins were removed by washing three times with PBST, and the protein complex was analyzed by immunoblotting. For Star-PAP-HSP27- PI4,5P_2_/PI3,4,5P_3_ binding assays, PI4,5P_2_ diC16 (#P-4516, Echelon Biosciences) and PI3,4,5P_3_ diC16 (#P-3916, Echelon Biosciences) were purchased. The phosphoinositide-mediated binding assay was performed in PBS by incubating constant amount of Star-PAP and HSP27 with an increasing amount of PI4,5P_2_/PI3,4,5P_3_ in the presence of 4 μg of Star-PAP antibody and 20 μl of nProtein A Sepharose. The other processes are as described above. Statistical data analysis was performed with Microsoft Excel, using data from at least three independent experiments.

### Microscale Thermophoresis (MST) Assay

MST assay was applied to measure the binding affinity of purified recombinant proteins *in vitro.* The target protein Star-PAP was fluorescently labeled by Monolith Protein Labeling Kit RED-NHS 2nd Generation (#MO-L011, Nano Temper) following the manufacturer’s instructions. A sequential titration of unlabeled ligand proteins, PI-PolyPIPosomes, or PI-micelles was made in a Tris-based MST buffer (50 mM Tris-HCl, pH 8.0, 50 mM NaCl, 80 mM KCl, 0.05% Tween-20) and mixed with an equal volume of fluorescently-labeled target protein prepared at 10 nM concentration in the same MST buffer, making the final target protein at a constant concentration of 5 nM and the ligand protein or lipid at a gradient. The effects of phoshoinositides were examined by incubating labeled Star-PAP with 2 μM PI-micelles before adding to the ligand titration describe above. The PI-PolyPIPosomes for MST were purchased from Echelon Biosciences, including PI PolyPIPosomes (#Y-P000, Echelon Biosciences), PI4,5P_2_ PolyPIPosomes (#Y-P045, Echelon Biosciences), and PI3,4,5P_3_ PolyPIPosomes (#Y-P039, Echelon Biosciences). The synthetic PIs for MST were also purchased from Echelon Biosciences, including PI diC16 (#P-0016, Echelon Biosciences), PI4,5P_2_ diC16 (#P-4516, Echelon Biosciences), and PI3,4,5P_3_ diC16 (#P-3916, Echelon Biosciences), which were dissolved in the MST buffer with 5 min sonication to prepare the PI-micelles. The target-ligand mixtures were loaded into Monolith NT.115 Series capillaries (#MO-K022, Nano Temper). The MST traces were measured by Monolith NT.115 pico, and the binding affinity was auto-generated by MO. Control v1.6 software.

### ^3^H-myo-inositol Metabolic Labeling

Cells were cultured in low serum Opti-MEM (#31985070, Thermo) supplemented with 10% dialyzed FBS with a 10,000 mw cut-off (#F0392, Sigma) and 1% Pen/strep (#15140-122, Gibco). In 10 cm dishes, 0.5 x 10^6^ HEK293FT cells were seeded to achieve ∼5% confluency the following day. 24 h after plating, cells were treated with either 1.322 μM (25 μCi/mL) of ^3^H-myo-inositol (#NET1156005MC, PerkinElmer) or unlabeled myo-inositol (#J60828.22, Thermo) and allowed to uptake the metabolic label for 48 h. 96 h after plating, cells were treated with control vehicle or 0.6% DEM for 4 h. For HA-Star-PAP overexpression, empty plasmid or HA-Star-PAP plasmid were transfected into the cells by branched-PEI 48 h after plating. Cells were then lysed and Star-PAP or HA-Star-PAP was immunoprecipitated before performing SDS-PAGE and CHCl3/MeOH extraction as indicated. SDS-PAGE samples were excised from the gel and dissolved in 30% H2O2 before liquid scintillation counting (LSC). Dissolved gel samples or resuspended protein samples were added to LSC vials with LSC cocktail (#6013319, PerkinElmer) before being processed by PerkinElmer Tri-Carb 4910 TR liquid scintillation analyzer. Analysis and DPM calculation were automated using QuantaSmart software.

## Statistics and Reproducibility

Two-tailed one-sample t-tests were used for pair-wise significance unless otherwise indicated. We note that no power calculations were used. Sample sizes were determined based on previously published experiments where significant differences were observed (Chen *et al*., 2022; Choi *et al*., 2019). Each experiment was repeated at least three times independently, and the number of repeats is defined in each figure legend. We used at least three independent experiments or biologically independent samples for statistical analysis.

## Acknowledgement

This work was supported in part by a National Institutes of Health grant R35GM134955 (R.A.A.), Department of Defense Breast Cancer Research Program grants HT9425-23- 1-0553 (V.L.C.), HT9425-23-1-0554 (R.A.A.) and W81XWH-21-1-0129 (V.L.C.), and a grant from the Breast Cancer Research Foundation (V.L.C.).

## Author Contribution

T.W., M.C, V.L.C., and R.A.A. designed the experiments. T.W. and M.C. performed the experiments. T.W., M.C., V.L.C., and R.A.A. wrote the manuscript.

## Data Availability

This study includes no data deposited in external repositories.

---

**Extended Data Figure 1.**
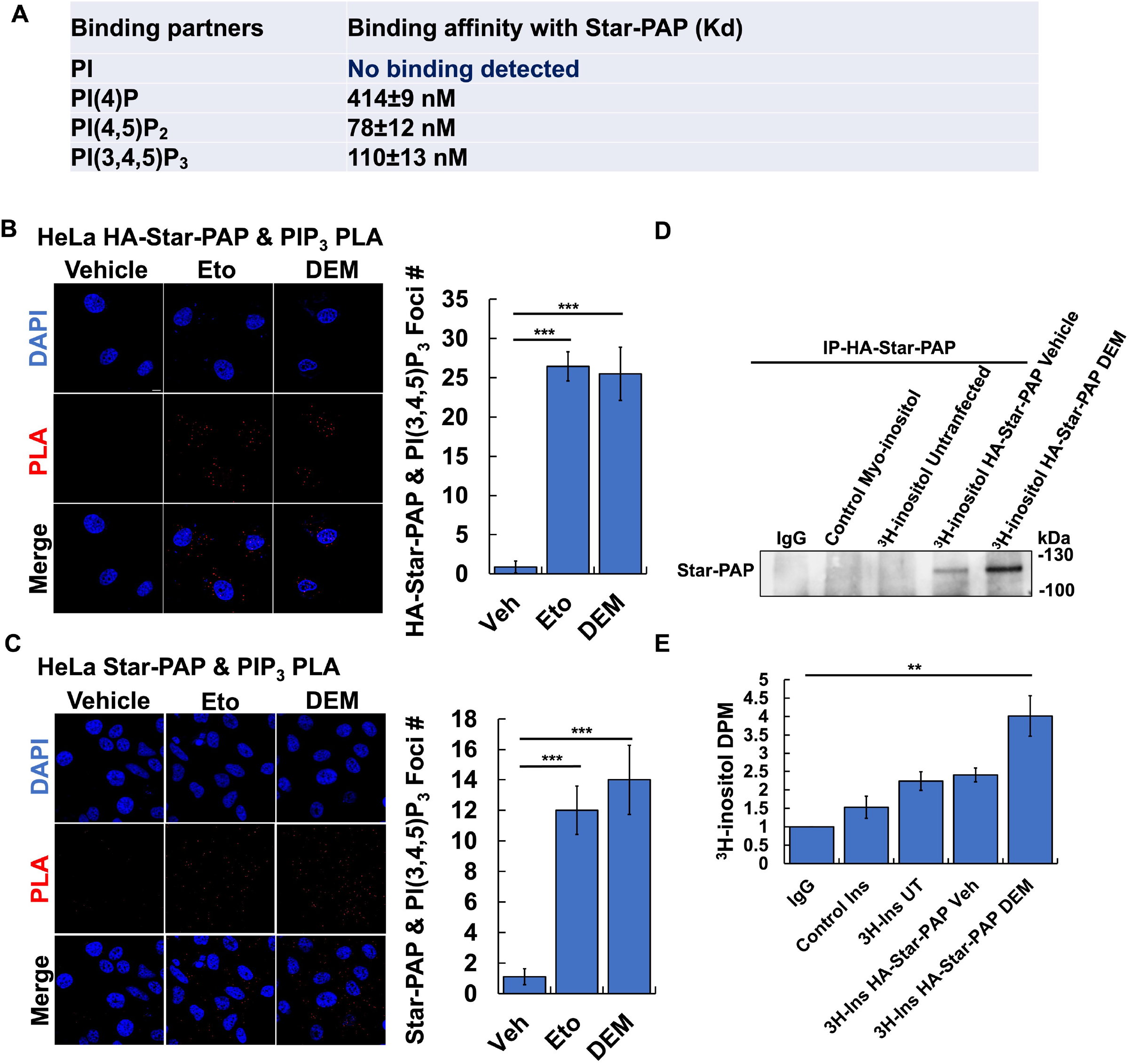
PI(4)P, PI(4,5)P_2_, and PI(3,4,5)P_3_ bind Star-PAP. A. The binding affinity of recombinant fluorescently labelled Star-PAP and non-labelled PIP_n_s was quantitated by MST assay. A constant concentration of fluorescently labelled Star-PAP (target, 5 nM) was incubated with increasing concentrations of non-labelled ligands to calculate the binding affinity. The binding affinities determined by MST are shown as indicated *K*d values. MST was performed using a Monolith NT.115 pico, and the binding affinity was autogenerated by MO. Control v.1.6 software. Data are presented as the mean ± s.d; *n* = 3 independent experiments. B. HeLa cells were transiently transfected with HA-tagged Star-PAP. 48 h later, cells were treated with 100 μM etoposide or 0.6% DEM for 4 and then processed for PLA analysis of HA and PI(3,4,5)P_3_ (*n* = 3 images each from three independent experiments, Scale bar, 10 μM). PLA foci per cell were quantified using ImageJ (*n* = 30 cells from three independent experiments). *P*-values are for one-sample *t*-test versus vehicle, *** for *P* < 0.001. C. HeLa cells were treated with 100 μM etoposide or 0.6% DEM for 4 h and then processed for PLA of Star-PAP and PI(3,4,5)P_3_ (*n* = 3 images each from three independent experiments, Scale bar, 10 μM). PLA foci per cell were quantified using ImageJ (*n* = 30 cells from three independent experiments). *P*-values are for one-sample *t*-test versus vehicle, *** for *P* < 0.001. **D, E.** HEK293FT cells were cultured from low confluency in media containing ^3^H myo-inositol or unlabeled control myo-inositol. After 24 h of growth cells were transfected with HA-Star-PAP plasmids. After 48 h of transfection, cells were treated with DEM for 4 h before being processed for IP against HA-tag. Star-PAP level was confirmed by WB (G) before samples were resolved by SDS-PAGE and the gel lane was excised and sectioned. Gel sections were then dissolved and analyzed by LSC, DPM were normalized to the IPed Star-PAP level (H). (n=3 independent experiments) *P*-values are for one-sample *t*-test versus vehicle, ** for *P* < 0.01.

**Extended Data Figure 2.**
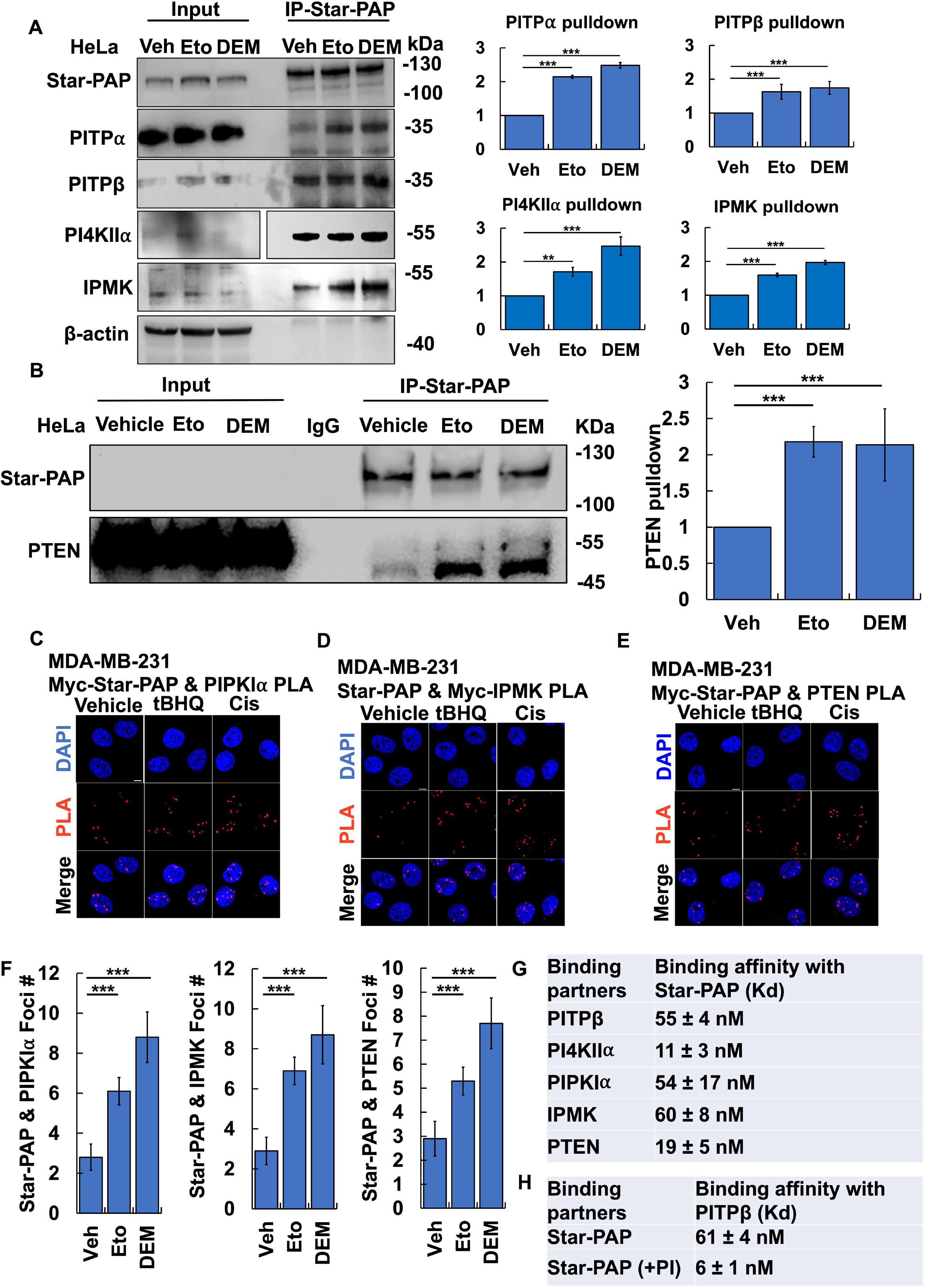
PITPs, PI kinases and phosphatases bind Star-PAP. A, B. HeLa cells were treated with 100 μM etoposide or 0.6% DEM for 4 h and processed for IP of Star-PAP and WB using antibodies against PITP⍺, PITPβ, PI4KII⍺, IPMK (A) and PTEN (B). PITP⍺/β and PTEN protein levels were quantified and normalized to the actin level (*n* = 3 independent experiments). *P*-values are for one-sample *t*-test versus vehicle, *** for *P* < 0.001. **C-E.** PLA images were performed using SP8 Leica confocal microscope, 100x with oil on MDA-MB-231 cells were transiently transfected with Myc-IPMK or Myc-Star-PAP. 48 h later, cells were treated with 1 mM tBHQ for 4 h or 30 μM Cisplatin for 24 h and then processed for PLA of Star-PAP and transfected IPMK or transfected Star-PAP and PIPKI⍺ or PTEN (*n* = 9 images from three independent experiments, Scale bar, 10 μM). **F.** PLA foci per cell in C-E were quantified using ImageJ (*n* = 10 cells from three independent experiments). *P*-values are for one-sample *t*-test versus control siRNA, ** for *P* < 0.01. **G.** The binding affinity of PITPβ, PI4KII⍺, PIPKI⍺, IPMK, and PTEN with Star-PAP was determined by MST. A constant concentration of fluorescently labelled Star-PAP (target, 5 nM) was incubated with increasing concentrations of non-labelled ligands to calculate the binding affinity. The binding affinities determined by MST are shown as indicated *K*d values. MST was performed using a Monolith NT.115 pico, and the binding affinity was autogenerated by MO. Control v.1.6 software. Data are presented as the mean ± s.d; *n* = 3 independent experiments. **H.** The binding affinity of Star-PAP with PITPβ in the presence and absences of phosphatidylinositol (PI) was determined by MST. A constant concentration of fluorescently labelled Star-PAP (target, 5 nM) was incubated with increasing concentrations of non-labelled ligands to calculate the binding affinity. For PI experiments, 2 μM of PI was incubated with labelled PITPβ before adding non-labelled Star-PAP. The binding affinities determined by MST are shown as indicated *K*d values. MST was performed using a Monolith NT.115 pico, and the binding affinity was autogenerated by MO. Control v.1.6 software. Data are presented as the mean ± s.d; *n* = 3 independent experiments.

**Extended Data Figure 4.**
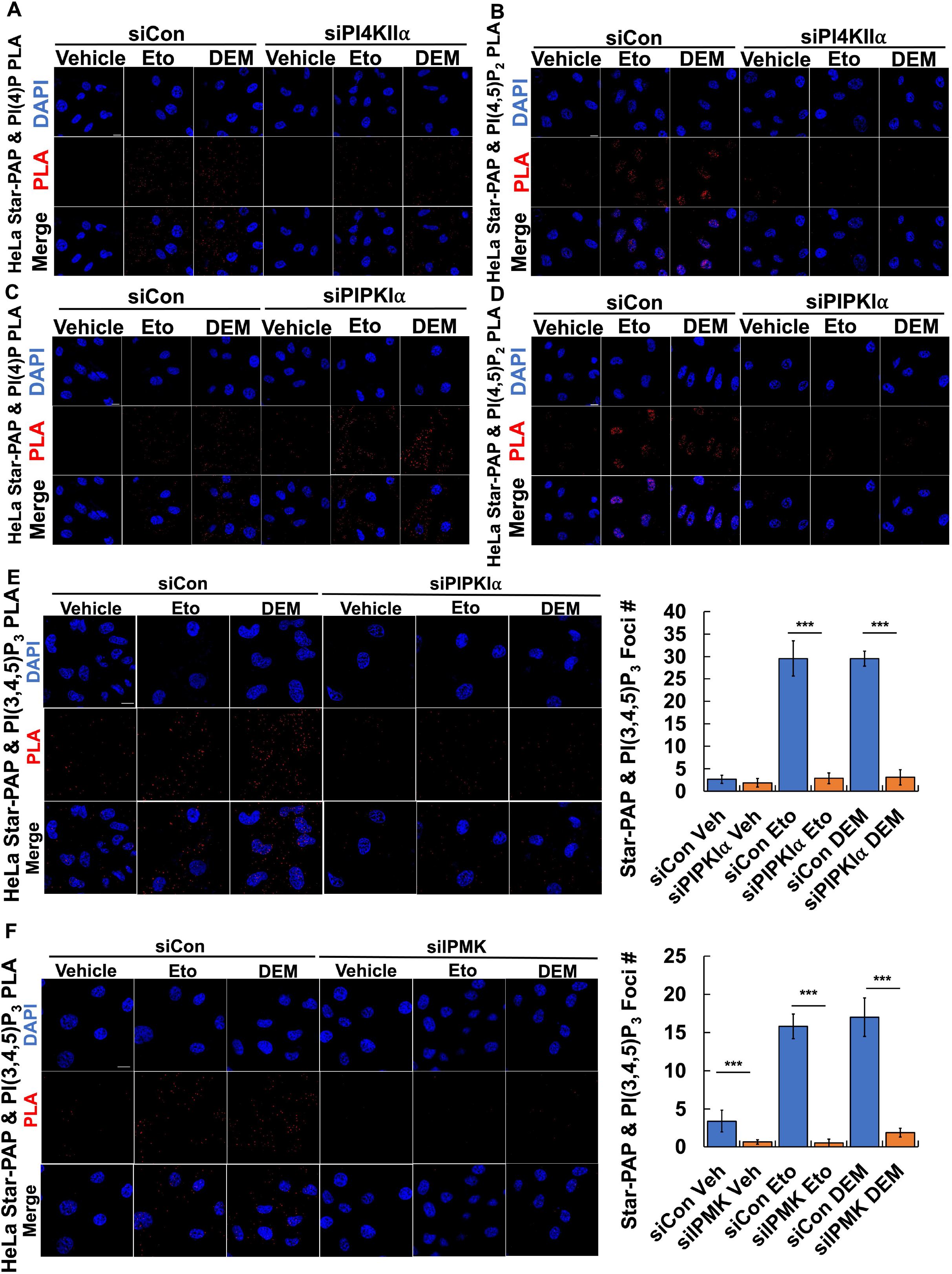
PI kinases regulate the interconversion of Star-PAP- coupled PIP_n_s. A, B. HeLa cells were transfected with control siRNAs or siRNAs targeting PI4KII⍺. 48 h later, cells were treated with 100 μM etoposide or 0.6% DEM for 4 h and then processed for PLA of Star-PAP and PI(4)P or PI(4,5)P_2_ (*n* = 3 images each from three independent experiments, Scale bar, 10 μM). **C, D.** HeLa cells were transfected with control siRNAs or siRNAs targeting PIPKI⍺. 48 h later, cells were treated with 100 μM etoposide or 0.6% DEM for 4 h and then processed for PLA of Star-PAP and PI(4)P or PI(4,5)P_2_ (*n* = 3 images each from three independent experiments, Scale bar, 10 μM). **E.** HeLa cells were transfected with control siRNAs or siRNAs targeting IPMK. 48 h later, cells were treated with 100 μM etoposide or 0.6% DEM for 4 h and then processed for PLA of Star-PAP and PI(3,4,5)P_3_ (*n* = 3 images each from three independent experiments, Scale bar, 10 μM). PLA foci per cell were quantified using ImageJ (*n* = 30 cells from three independent experiments). *P*-values are for one-sample *t*-test versus vehicle, ** for *P* < 0.01, *** for *P* < 0.001. **F.** HeLa cells were transfected with control siRNAs or siRNAs targeting PIPKI⍺. 48 h later, cells were treated with 100 μM etoposide or 0.6% DEM for 4 h and then processed for PLA of Star-PAP and PI(3,4,5)P_3_ (*n* = 3 images each from three independent experiments, Scale bar, 10 μM). PLA foci per cell were quantified on ImageJ (*n* = 30 cells from three independent experiments). *P*-values are for one-sample *t*-test versus vehicle, ** for *P* < 0.01, *** for *P* < 0.001.

**Extended Data Figure 5.**
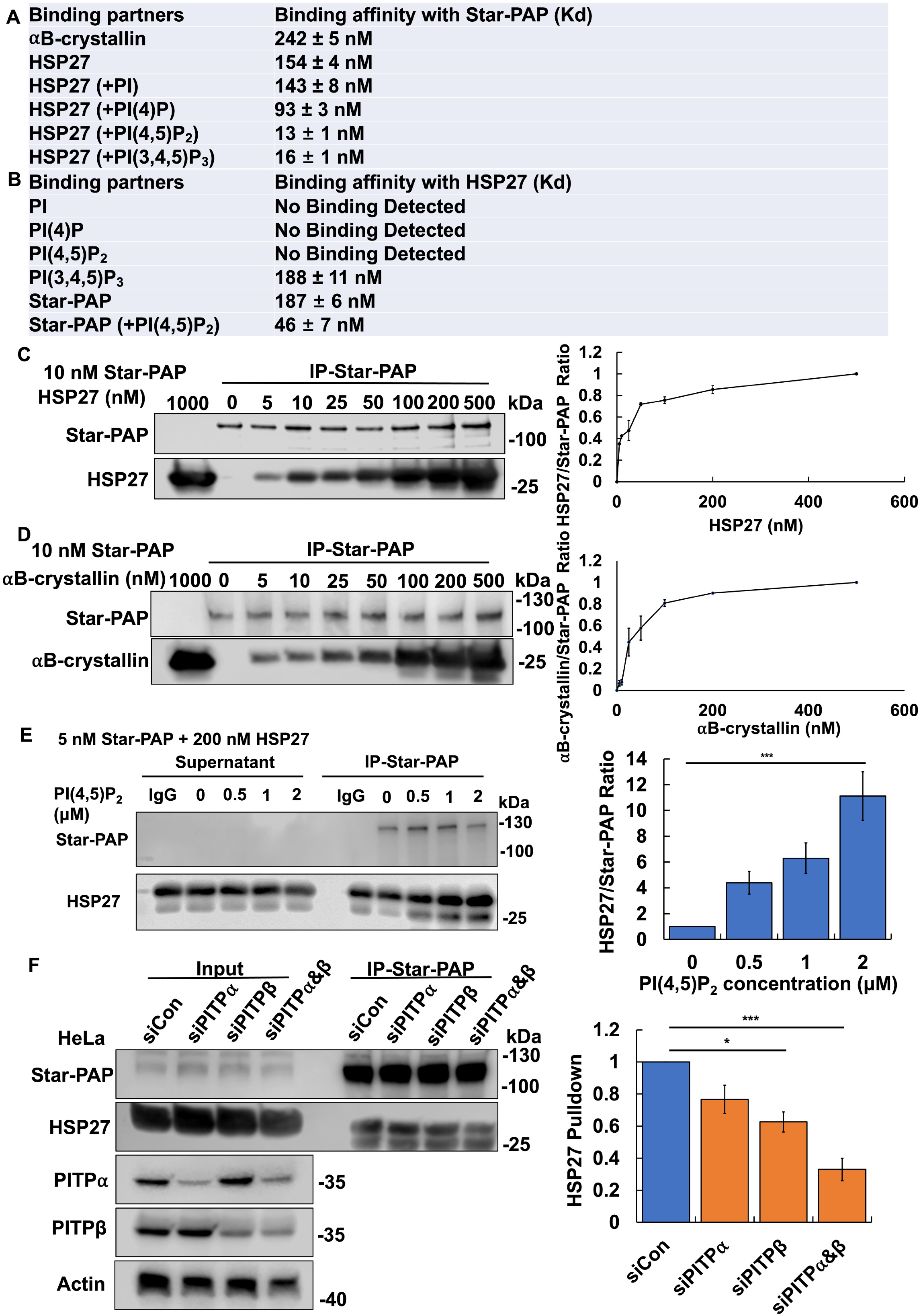
sHSPs bind Star-PAP by a PIPn-dependent mechanism. **A.** The binding affinity of HSP27 and ⍺B-crystallin with Star-PAP, and the effects of PIP_n_s on these interactions was determined by MST. A constant concentration of fluorescently labelled Star-PAP (target, 5 nM) was incubated with increasing concentrations of non-labelled ligands to calculate the binding affinity. In some experiments, 2 μM of PI, PI(4)P, PI(4,5)P_2_ or PI(3,4,5)P_3_ were incubated with labelled Star-PAP before adding non-labelled HSP27. The binding affinities determined by MST are shown as indicated *K*d values. MST was performed using a Monolith NT.115 pico, and the binding affinity was autogenerated by MO. Control v.1.6 software. Data are presented as the mean ± s.d; *n* = 3 independent experiments. **B.** The binding affinity of PIP_n_s and Star-PAP with HSP27, and the effects of PI(4,5)P_2_ on the interaction of Star-PAP and HSP27 were examined by MST. A constant concentration of fluorescently labelled HSP27 (target, 5 nM) was incubated with increasing concentrations of non-labelled ligands to calculate the binding affinity. In some experiments, 2 μM of PI(4,5)P_2_ were incubated with labelled Star-PAP before adding non-labelled HSP27. The binding affinities determined by MST are shown as indicated *K*d values. MST was performed as described in A. **C,D.** *In vitro* binding assay was performed by adding increasing amount of His-tagged HSP27 protein (C) or His-tagged ⍺B-crystallin protein (D) to 10 nM His-tagged Star-PAP protein. Star-PAP was then IPed and WB for Star-PAP and ⍺B-crystallin was performed (*n* = 3 independent experiments). Protein levels were quantified using an Odyssey Imaging System (LI-COR Biosciences). The binding curve is drawn through the HSP27/Star-PAP or ⍺B-crystallin/Star-PAP signal ratio of each condition. **E.** *In vitro* binding assay was performed by adding increasing amounts of PI(4,5)P_2_ to 5 nM Star-PAP and 200 nM HSP27. Star-PAP was then IPed and WB for Star-PAP and HSP27 was performed (*n* = 3 independent experiments). HSP27 levels were quantified and normalized to the Star-PAP level of each condition. *P*-values are for one-sample *t*- test versus 0 μM PIP_2_, *** for *P* < 0.001. **F.** HeLa cells were transfected with control siRNAs or siRNAs targeting PITP⍺, PITPβ, or both PITP⍺/β . 48 h later, cells were processed for Star-PAP IP followed by WB for Star-PAP and HSP27 (*n* = 3 independent experiments). HSP27 level were quantified and normalized to the Star-PAP level in each condition. *P*-values are for one-sample *t*-test versus control siRNA, * for *P* < 0.05, *** for *P* < 0.001.

**Extended Data Figure 6.**
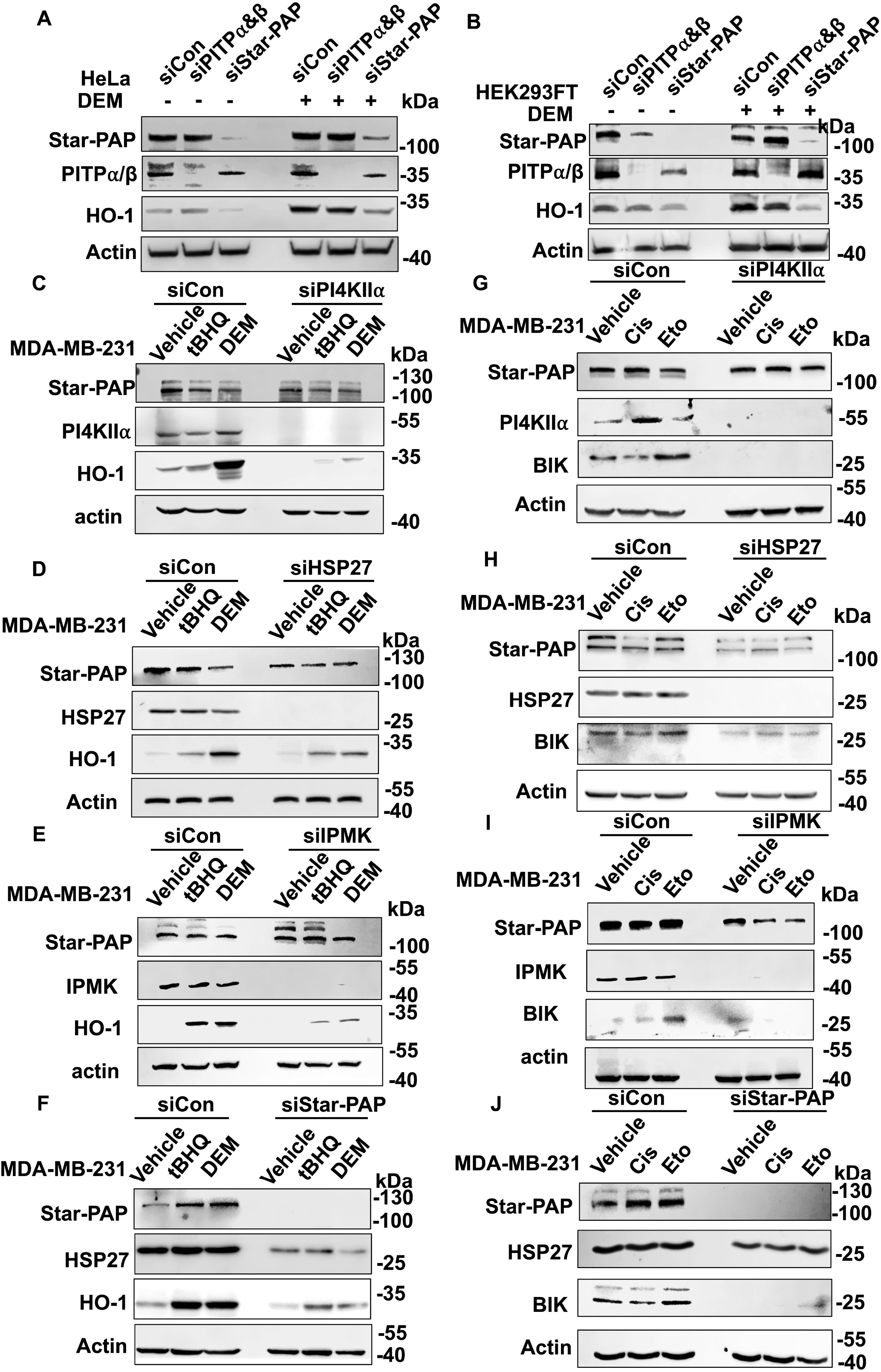
WB images of the effects of PITP⍺/β, HSP27, IPMK and Star-PAP KD on HO-1 and BIK levels. A. HeLa cells were transfected with control siRNAs or siRNAs targeting both PITP⍺/β or Star-PAP. 48 h later, cells were treated with 0.6% DEM for 4 h and analyzed by WB for HO-1 (*n* = 3 independent experiments). **B.** HEK293FT cells were transfected with control siRNAs or siRNAs targeting both PITP⍺/β or Star-PAP. 48 h later, cells were treated with 0.6% DEM for 4 h and analyzed by WB for HO-1 (*n* = 3 independent experiments). **C-F.** MDA-MB-231 cells were transfected with control siRNAs or siRNAs targeting PI4KII⍺ knockdown (C), HSP27 knockdown (D), IPMK knockdown (E), or Star-PAP knockdown (F). 48 h later, cells were treated with 1 mM tBHQ or 0.6% DEM for 4 h and analyzed by WB for HO-1 (*n* = 3 independent experiments). **G-J.** MDA-MB-231 cells were transfected with control siRNAs or siRNAs targeting PI4KII⍺ knockdown (G), HSP27 knockdown (H), IPMK knockdown (I), or Star-PAP knockdown (J). 24 h later, cells were treated with 30 μM Cisplatin for 24 h or 100 μM etoposide for 4 h and analyzed by WB for BIK (*n* = 3 independent experiments).

